# Kidney cortex macrophages strategically reabsorb phosphate from the urine to prevent the formation of mineral stones

**DOI:** 10.1101/2025.02.18.638798

**Authors:** Yuxi Wang, Yuancheng Weng, Xukai Ding, Ningting Chen, Qiang Wang, Zelin Lu, Ziqi Gao, Jian He, Feng Xu, Dandan Liang, Fei Han, Xiao Z Shen

## Abstract

Resident macrophages are important in maintaining tissue homeostasis by meeting specialized physiological demands and mitigating the unique stresses each tissue endures. In the kidney, urine is formed in a process of glomerular filtration and tubular reabsorption. Supersaturation of mineral solutes such as calcium phosphate poses a persistent challenge to the kidney; if left unchecked, kidneys stones form, which will cause tubular obstruction and even kidney failure. However, cellular mechanisms of preventing kidney stone formation remain incompletely understood. In this study, we found that resident macrophages in kidney cortex distinguishably expressed phosphate transporter SLC34A1. These cells extend transtubular protrusions and were capable of reabsorbing phosphate from the urine through the SLC34A1, a function previously only attributed to tubular epithelial cells. Depletion of *Slc34a1* from kidney cortex macrophages led to increased urinary phosphate excretion, disrupting body phosphorus balance. More importantly, loss of macrophage-mediated phosphate reabsorption from the urine resulted in a drastic over-deposition of calcium phosphate microcrystals in kidney tubules, particularly in the late proximal tubules of inner cortex where intratubular calcium concentration is high. This also accelerated mineral stone formation. Exposure to mineral crystals stimulated cortex macrophages to upregulate SLC34A1 expression, a response mechanistically driven by lysosomal disruption. As such, cortex macrophages employed phosphate reabsorption as a proactive strategy to prevent mineral stone formation.

## Introduction

The kidney consists of cortex and medulla. Its tubular system extends from the corpuscle exclusively distributed in the cortex, collects glomerular filtrates and drains urine to the collecting duct in the medulla, and eventually empties into the renal pelvis. Although the physicochemical property of filtration barrier in kidney corpuscle precludes the passage of large blood-borne particles, e.g., blood cells, into the urine, the urine usually contains a plethora of particles derived from varying sources. The most pronounced particles are mineral crystals which when sedimentating and aggregating, would form kidney stones and block urine passage. The formation of mineral stones is due to supersaturation(*1*), as ∼99% of water from the glomerular filtrate is reabsorbed during its passage through renal tubules(*2*). Reabsorption of mineral solutes from the urine occur in the kidney as well to avoid unnecessary waste, but reabsorption degrees vary for different electrolytes. Electrolyte reabsorption ability is so far exclusively attributed to tubular epithelial cells, especially those lining the proximal tubules in the cortex(*3*). The degrees of urinary supersaturation causally affect nephrolithiasis (formation of kidney stone) and up to a quarter of urine samples from healthy subjects show crystalline particles(*4*).

During nephrolithiasis, once a crystal nucleus is established inside the kidney, exposure to the urine minerals enables the stone to grow within the renal tubules(*5*). Globally, hydroxyapatite, a basic calcium phosphate (CaPi) crystal, is the second most common stone type after calcium oxalate (CaOx)(*6*). However, approximately 80% of kidney stones are composed of CaOx mixed with CaPi(*7*), and CaPi by itself is also a good nucleator of CaOx(*8*) and has been shown to transform itself into CaOx through dissolution and recrystallization(*9, 10*). In the kidney, there is no known oxalate reabsorption transporter, and oxalate filtrate plus secretion of oxalate in the tubules results in absolute oxalate excretion(*11*). In contrast, healthy organisms are protected from urinary phosphorus loss by constitutively active tubular phosphate (Pi) reabsorption, which is mainly mediated by the sodium-phosphate cotransporters SLC34A1 expressed in the proximal tubules(*12, 13*). Mutations in SLC34A1 cause nephrocalcinosis and kidney stones in human(*14, 15*). Regarding the central cation constituting the kidney stones, Ca^2+^ is predominantly reabsorbed in the late proximal tubules via passive paracellular transport based on transepithelial concentration gradient(*16*). Although the etiology of nephrolith have been extensively studied(*6, 17–19*), the cellular mechanisms underlying the protection from sedimentary particle formation/aggregation just emerged. We recently reported that the resident macrophages (MØ) positioned in the close proximity to tubules constitutively formed transtubular protrusions and these behaviors were present in both cortex and medulla(*20*). The medulla MØ preferentially removed intratubular preformed sedimentary particles including mineral crystals to prevent blockade(*20*). What remains unclear is the role of cortex juxtatubular MØ and whether there is an inherent mechanism to prevent the supersaturation of mineral solutes in the urine so as to limit the formation of mineral crystals.

In this study, we revealed that distinctive from resident MØ in other tissues, kidney cortex MØ highly express the Pi transporter SLC34A1 and could reabsorb Pi from the urine, which is integral to maintain the phosphorus homeostasis of body. More importantly, the cortex MØ would upregulate SLC34A1when sensing the presence of mineral microcrystals in the tubules, as a strategy to decrease CaPi supersaturation and limit kidney stone formation by reabsorbing urine Pi.

## Results

### Kidney cortex macrophages distinctively express phosphate transporter SLC34A1

The cortex and medulla of kidney are different in cellular composition, structure, function, and microenvironment(*2*). MØ distribute in both cortex and medulla in steady state, and our previous work revealed that MØ in these anatomic areas differ in transcriptome (*20*), implying that they are affected by different environmental cues and play different homeostasis-maintaining roles. Among the over 2,000 differentially expressed genes (DEGs, defined by fold change > 2; adjusted P value < 0.001) between cortex and medulla MØ(*20*), gene ontology (GO) enrichment for biological process analysis showed that the cortex MØ were characterized by enriched expression of transcripts involved in anion transport (Fig. 1A). In particular, *Slc34a1*, which encodes sodium-dependent phosphate transporter 2A (NaPi-IIa) previously thought to be exclusively expressed by renal proximal tubules(*21, 22*), was notably expressed in the cortex MØ among the 95 anion-transport-associated DEGs upregulated in the cortex MØ (Fig. 1B). Ranking all known phosphate transporter genes based on expression levels disclosed that *Slc34a1* was the most abundant phosphate transporter transcribed in cortex MØ (among the top 4% highly expressed genes), while *Slc34a1* expression in the medulla MØ was negligible (Fig. 1C). This distinctive expression patterns of *Slc34a1* between cortex and medulla MØ were verified in *Slc34a1*^CreERT2/+^*:R26*^tdTomato^ reporter mice when examined by flow cytometry (Fig. 1D). To be noted, upon flow cytometry analysis, we particularly precluded AQP1^+^ cells (fig. S1) to avoid contamination from proximal epithelial cells. Our previous work showed that the majority (over 60%) of cortex MØ were in close proximity to kidney tubules, and they could form transtubular protrusions exposed to tubular lumen (fig. S2)(*20*). Histological study confirmed that the MØ attached to the proximal tubules express tdTomato in addition to the proximal tubules in the SLC34A1 reporter mice (Fig. 1E).

**Fig. 1.**
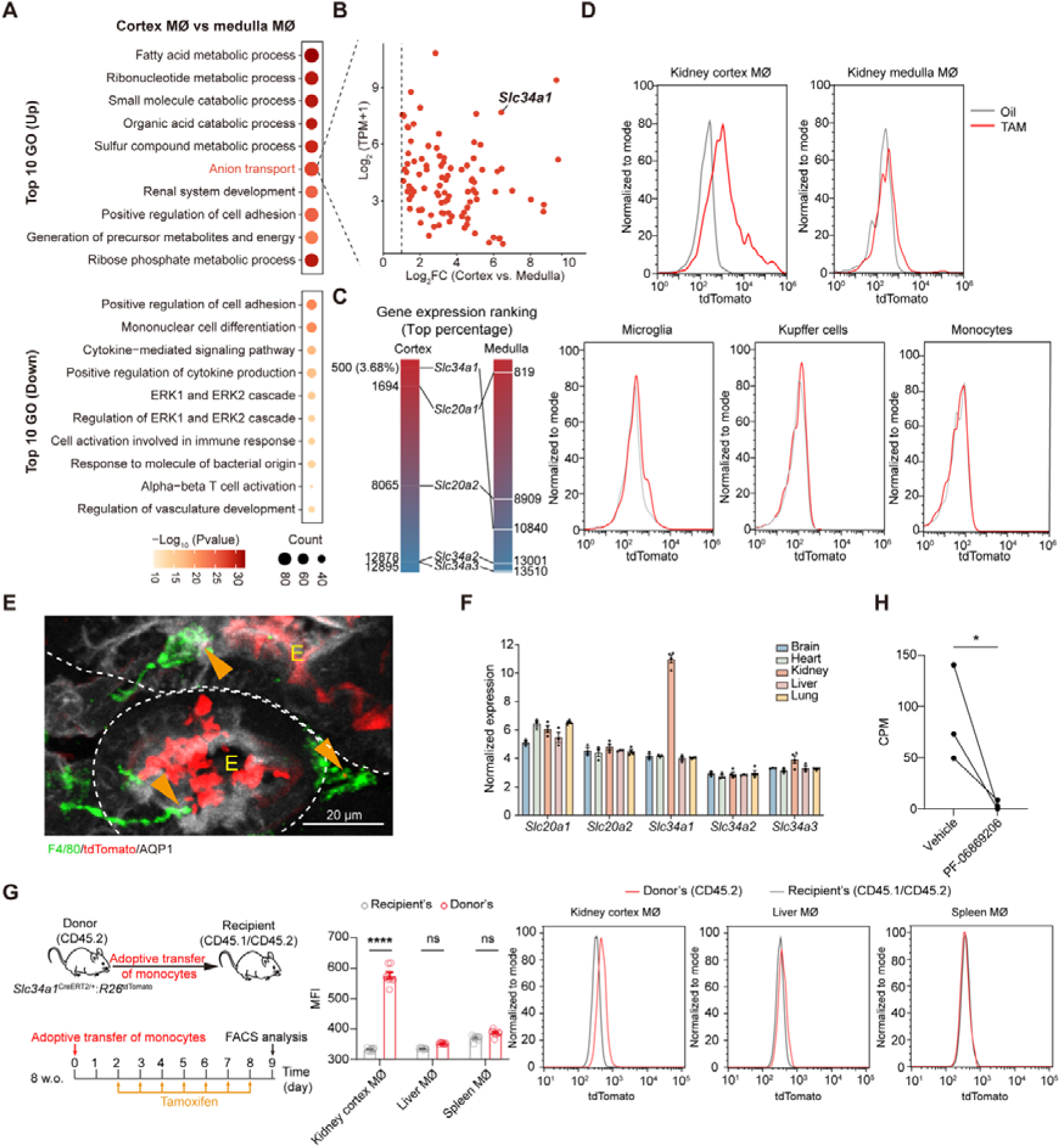
Kidney cortex macrophages distinctively express SLC34A1. (**A**) Top 10 upregulated and downregulated pathways of cortex MØ relative to medulla MØ, derived from GO analysis of DEGs. (**B**) Expression levels and ratios of the DEGs in the “Anion transport” term of GO analysis. *Slc34a1* is highlighted. (**C**) The average rankings regarding the expression abundance of all Pi transporters in the cortex and medulla MØ according to gene expression values (TPM). (**D**) Relative expression of SLC34A1 in the kidney cortex and medulla MØ, microglia, Kupffer cells and blood monocytes of the *Slc34a1*^CreERT2/+^*:R26*^tdTomato^ mice, as indicated by tdTomato expression. (**E**) Representative confocal images of cortex sections show the expression of SLC34A1 (tdTomato) in juxtatubular MØ of *Slc34a1*^CreERT2/+^*:R26*^tdTomato^ reporter mice. Dotted line: the basolateral border of tubule. E, epithelial cells. Arrowhead, tdTomato expression in MØ. (**F**) The relative expression levels of all Pi transporter genes in resident MØ derived from the indicated organs, which was calculated by averaging the normalized gene expression according to the normalized data using the robust multi-array average by the authors(*23*). (**G**) Monocytes derived from the bone marrow of *Slc34a1*^CreERT2/+^*:R26*^tdTomato^ mice were adoptively transferred *i.v.* into naive mice, and their fates were traced in the kidney, liver and spleen. The representative expression patterns of tdTomato in donor-derived MØ in these organs were shown and their mean fluorescent intensities (MFI) were compared with those of recipients’ MØ in the same organs as tdTomato^−^ controls. (**H**) Kidney cortex MØ were purified and pre-treated with PF-06869206, a SLC34A1-specific blocker, or vehicle, followed by being incubated with ^32^P-labeled phosphate solution for 15 min. The intracellular ^32^P radioactivity was then measured. Each dot represents a pool of cells derived from 3 mice. ns, not significant. \**P*<0.05, \*\*\*\**P*<0.001 by two-way ANOVA (G) and one-tailed paired *t* test (H). Data are depicted as mean±SEM. Data in D, E, G and H are derived from at least 3 independent experiments.

Actually, expression of *Slc34a1* is a characteristic of kidney cortex MØ since data mining of a transcriptomic study of MØ derived from multiple organs(*23*) revealed that kidney MØ distinguishably expressed a high level of *Slc34a1* relative to other tissue MØ (Fig. 1F). Using the *Slc34a1* reporter mice, we confirmed that in contrast to kidney cortex MØ, resident MØ of the brain (microglia) and liver (Kupffer cells) barely expressed SLC34A1 (Fig. 1D and fig. S3A). We previously showed that most of kidney cortex MØ were postnatally monocyte-derived, while medulla MØ were embryo-derived and self-maintained in adult mice(*20*). However, blood monocytes did not have an obvious tdTomato expression in the *Slc34a1* reporter mice (Fig. 1D), implying that SLC34A1 expression occurs when monocytes infiltrate and differentiate to MØ in the kidney. To probe this hypothesis, we performed an adoptive transfer strategy by *i.v.* infusing monocytes derived from *Slc34a1*^CreERT2/+^:*R26*^tdTomato^ mice (in a CD45.2 background) into CD45.1/CD45.2 hybrid mice (Fig. 1G). Previously, we showed that some *i.v.* transferred monocytes would arrive in the kidneys and differentiated to MØ by over a week(*24*). Starting from day 2 post transfer, the recipients were treated with tamoxifen for a week. We noticed that on day 9 post transfer, a significant portion of donor-derived MØ in the kidney cortex expressed tdTomato, which was not observed in the liver and spleen of the same recipients (Fig. 1G and fig. S3B). These data indicate that expressing SLC34A1 is a unique character of MØ in the kidney cortex, possibly impacted by the special kidney niche they reside in. To ascertain that the SLC34A1 expressed by cortex MØ is functional, we purified MØ from kidney cortex of C57BL/6 mice and validated no contamination from epithelial cells (fig. S4A). The cortex MØ were pulsed with ^32^P-labeled Pi in culture. By application of PF-06869206(*25*), a SLC34A1-specific blocker, we demonstrated that these cells could uptake extracellular Pi in a SLC34A1-dependent manner (Fig. 1H).

### Kidney cortex macrophages routinely reabsorb phosphate from the urine

Phosphate reabsorption mainly takes place in the proximal tubules and relies on epithelial SLC34A1. Kidney MØ homogenously highly express CX3CR1(*20*) and *Cx3cr1*^CreERT2/+^:Ai14 reporter mice exhibited that tubule epithelial cells did not express CX3CR1 (fig. S5). To interrogate whether cortex MØ could also participate in Pi reabsorption *via* SLC34A1, we crossed *Cx3cr1*^CreERT2/+^ mice with *Slc34a1*^fl/fl^ mice (Fig. 2A). Intraperitoneal tamoxifen treatment reduced *Slc34a1* transcripts by 79.3% in cortex MØ of the offspring (hereafter *Slc34a1*^ΔMØ^) (Fig. 2B). Abrogation of *Slc34a1* expression by cortex MØ neither appeared to alter their density or morphology (fig. S6A), nor stirred an upregulation of proinflammatory cytokines (fig. S6B). Indeed, an acute deprivation of MØ *Slc34a1* expression resulted in an increased urinary Pi excretion (hyperphosphaturia) (Fig. 2C). Not surprisingly, since SLC34A1 is a sodium-dependent phosphate transporter, the mice also had an increased Na^+^ output, while the excretion of both K^+^ and Cl^−^ in the urine were unaltered (Fig. 2C). These results were consistent when the values were adjusted by either urine creatinine (Fig. 2C) or body weight (fig. S6C). Over-excretion of Pi led to a decrease of parathyroid hormone (PTH) with an unaltered fibroblast growth factor 23 (FGF23) in blood plasma (Fig. 2D), consistent with the phenotypes of mice with a complete SLC34A1 depletion by either genetic or pharmacological approaches(*25, 26*). It was reported that Pi over-excretion from the urine would elicit an adaptive increase of calcitriol (Vitamin D_3_) production(*26*). Calcitriol production is determined by calcitriol anabolic and catabolic enzymes encoded by *Cyp27b1* and *Cyp24a1* in the kidney, respectively(*27, 28*). Over-excretion of Pi in the *Slc34a1*^ΔMØ^ mice was accompanied with an attendant elevation of *Cyp27b1* but not *Cyp24a1* transcripts in the kidney (Fig. 2E), indicating calcitriol overproduction. Due to the key regulation of calcium reabsorption by PTH, the decreases of PTH would ultimately result in hypercalciuria which was observed in the *Slc34a1*^ΔMØ^ mice (Fig. 2C), in alignment with the phenotype with a complete abrogation of SLC34A1 activity(*26*). PTH could suppress SLC34A1 expression in proximal epithelial cells(*29*). Under the influences of a decreased PTH in the *Slc34a1*^ΔMØ^ mice, *Slc34a1* expression in the cortex tubules was modestly upregulated (fig. S6D), which also suggests that the depleting effect on *Slc34a1* expression was not leaked to proximal tubules epithelial cells in *Slc34a1*^ΔMØ^ mice. The proximal tubules were generally intact since a normal structure and density of proximal tubules was shown (fig. S6E); the expression of *Aqp1* (the gene encoding Aquaporin 1) and *Slc9a3* (the gene encoding Na^+^/H^+^ exchanger 3), two signature transporters of proximal tubules, were not altered (fig. S6F); also, glucose excretion through urine was not elevated (fig. S6G), suggesting that tubulopathy was not responsible for the renal Pi leak in the mutant mice. Assessment of tissue integrity and inflammatory infiltrates by hematoxylin and eosin (H&E) staining revealed no histopathologic evidence of kidney inflammation in the mice deprived of MØ *Slc34a1* expression (fig. S6H). In addition, abrogation of *Slc34a1* expression in MØ neither affected global kidney functions, as evaluated by plasma creatinine and urea nitrogen (BUN) levels (fig. S6I), nor altered glomerulus function, as assessed by glomerulus filtration rate (GFR) (fig. S6J).

**Fig. 2.**
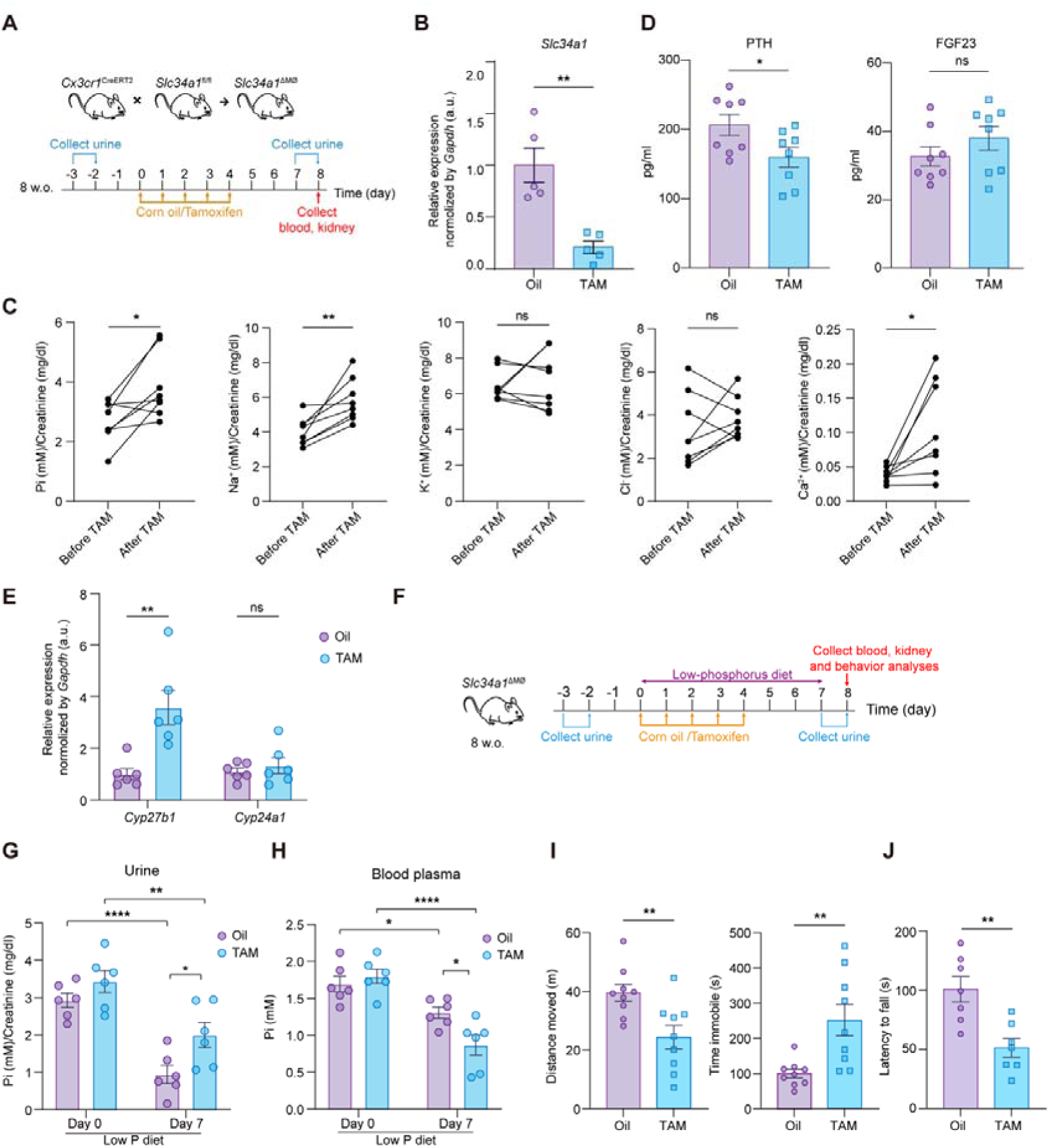
Kidney cortex macrophages constitutively reabsorb Pi from the urine. (**A**) Scheme illustrating the generation of *Slc34a1*^ΔMØ^ mice and the experimental protocol of Panels B-E in which *Slc34a1*^Δ MØ^ mice were treated with either corn oil (Oil) or tamoxifen (TAM). (**B**) Efficiency of *Slc34a1* deletion in kidney cortex MØ, examined by RT-PCR. (**C**) Urine output of the indicated electrolytes, adjusted by the excreted creatinine, before and after TAM treatment. (**D**) The concentrations of PTH and FGF23 in the blood plasma. (**E**) The relative expression of *Cyp27b1* and *Cyp24a1* transcripts in the kidney. (**F**) Scheme illustrating the protocol of low-phosphorus diet experiment for Panels G-J. (**G** and **H**) Urine Pi output (G) and Pi concentrations in blood plasma (H). (**I**) In open-field test, after the indicated treatments, *Slc34a1*^ΔMØ^ mice were placed in a square arena, and their total moving distance (left) and immobile time (right) were recorded during the observation period. (J) In rotarod test, after the indicated treatments, *Slc34a1*^ΔMØ^ mice were placed on a rod which then started to rotate in an accelerated speed. The on-rod time periods before their falling were recorded. ns, not significant. \**P*<0.05, \*\**P*<0.01, \*\*\*\**P*<0.001 by two-tailed unpaired *t* test (B, D, E, I, J), two-tailed paired *t* test (C) and two-way ANOVA (G, H). Data are depicted as mean±SEM. Data are derived from at least 2 independent experiments.

Administration of tamoxifen into the subcapsular space of kidney would limit its effect on the kidney(*30*), and *Slc34a1*^ΔMØ^ mice again showed hyperphosphaturia and hypercalciuria (fig. S7A). Moreover, C57BL/6 mice *i.p.* treated with tamoxifen did not develop hyperphosphaturia or hypercalciuria (fig. S7B). These experiments exclude an off-target effects induced by *i.p.* tamoxifen administration. Intravesical application of diphtheria toxin (DT) to *Cx3cr1*^CreERT2/+^:*iDTR* mice could specifically deplete kidney MØ(*20*), and we found that depleting kidney MØ could also lead to hyperphosphaturia and hypercalciuria (fig. S7C), substantiating our finding.

Albeit with an excessive excretion of Pi *via* the urine, *Slc34a1*^ΔMØ^ mice had no overt hypophosphatemia on sacrifice at the end of the experimental protocol (fig. S7D), possibly due to a compensation by an increased dietary phosphorus absorption, especially considering that an elevated calcitriol enhances phosphorus absorption in the small intestine(*31*). Thus, we next set to evaluate the importance of kidney MØ-dependent Pi reabsorption on health by feeding the *Slc34a1* ^Δ MØ^ mice with a low-phosphorus diet. To do that, tamoxifen- or vehicle-treated *Slc34a1*^ΔMØ^ mice were fed with a phosphorus-restricted diet (0.02% vs. 0.6-1.2% in the normal chow) for 1 wk (Fig. 2F). The efficacy of phosphorus intake restriction was confirmed by a significantly lower urinary Pi excretion in the control mice after 1 wk of diet (Fig. 2G). Although the output of Pi was also reduced in the mutated mice under the same diet, it was still higher than that of the controls (Fig. 2G). As such, a more profound reduction of Pi concentration in blood plasma occurred in the mutated mice (to 48.2% of baseline *vs.* to 77.1% of baseline in controls) (Fig. 2H). The severe hypophosphatemia in the mutated mice was accompanied with movement impairment and muscle weakness(*32*), as manifested by open-field test and rotarod test, respectively (Figs. 2I and 2J). These data together suggest kidney cortex MØ facilitate in maintaining phosphorus homeostasis by expressing SLC34A1, particularly when phosphorus intake is limited.

### SLC34A1 expression by kidney cortex MØ is critical in preventing kidney stone formation

Unsurprisingly, the *Slc34a1*^ΔMØ^ mice generally displayed a milder Pi-associated metabolic alteration in comparison to the previously reported *Slc34a1*-null mice under normal diet(*26*), e.g., milder increase of urine Pi excretion (40% *vs.* 150%), no difference *vs.* 35.2% decrease in blood serum Pi, and a milder PTH response (22.5% *vs.* 72.9% reduction), indicating a minor role of MØ in Pi reabsorption for body phosphorus homeostasis relative to that of tubule epithelial cells. However, considering that Pi is significantly concentrated in the urine (99% of water is reabsorbed, while ∼10-20% of Pi in the filtrate of the glomeruli is eventually excreted in terminal urine)(*33*), the loss of MØ’s ability in reabsorbing Pi may place a significant threat to an oversaturation of Pi and formation of CaPi crystals. We thus predicted that besides the increased excretion of Pi in the urine as a soluble form (Fig. 2C), there were more sedimentary CaPi crystals formed and deposited in the kidneys of the MØ *Slc34a1*-deficient mice as well. To probe this hypothesis, we imaged the kidneys employing a fluorescent bisphosphonate (OsteoSense) that can bind to crystalline CaPi and has a high detection sensitivity(*34*). Indeed, there was a remarkable 10-fold increase of OsteoSense^+^ crystals in the kidney, in particular in the inner cortex, of the MØ *Slc34a1*-ablated mice compared to the control mice (Fig. 3A). Urine acidification could reduce CaPi crystals. By giving an ammonium chloride solution as the drinking water to mice(*34*), we observed reduced OsteoSense^+^ microparticles in the tamoxifen-treated *Slc34a1*^ΔMØ^ (Fig. 3A), further verifying their CaPi identity. Of note, these minute CaPi particles could not be discerned by Alizarin Red S staining, a regular way for detecting kidney mineral stones.

**Fig. 3.**
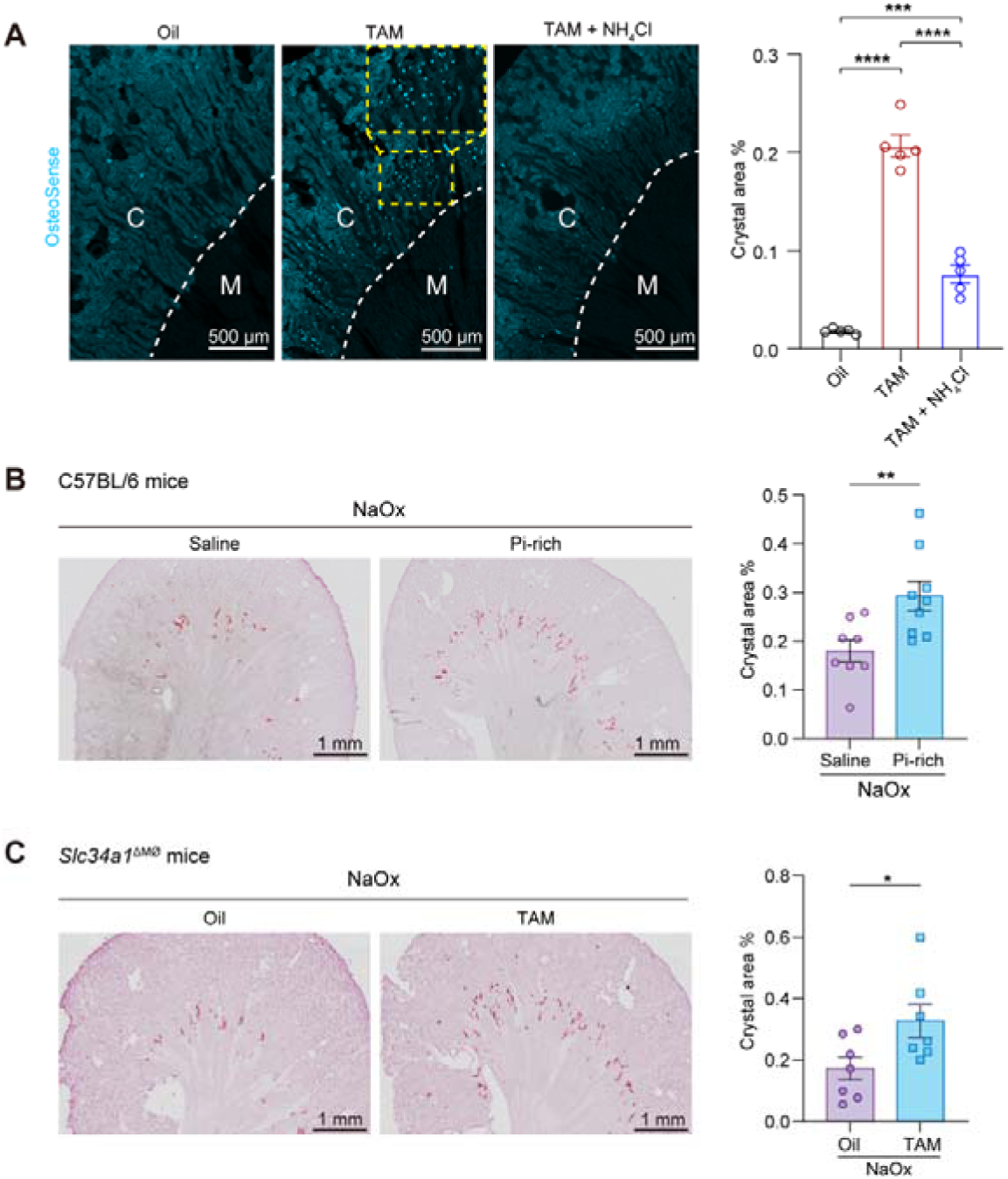
Loss of *Slc34a1* expression by kidney cortex macrophages results in a rampant CaPi crystal deposition in kidney tubules and an accelerated kidney stone formation. (**A**) *Slc34a1*^ΔMØ^ mice were treated with Oil, TAM or TAM plus NH_4_Cl. CaPi particles (blue dots) highlighted by OsteoSense detection were shown. CaPi crystal areas in the kidney were measured. White dotted line: border of cortex and medulla (kidney cortex has relatively high autofluorescence). Each dot represents the average of the affected areas of four whole-mount slides from one mouse. C, cortex. M, medulla. n = 5. (**B**) Alizarin red S staining of the kidneys of C57BL/6 mice after a NaOx *i.p.* regimen (60 mg/kg/day for 3 consecutive days). Some of the mice were *i.p.* co-treated with Pi-rich solution (Na_2_HPO_4_-NaH_2_PO_4_ with a final concentration of 100 mM phosphate, pH 7.2 at a dose of 100 μl per 25g body weight.). Each dot represents the average of the affected areas of 4 whole-mount slides from one mouse. (**C**) After the Oil or TAM treatment, *Slc34a1*^ΔMØ^ mice were administered *i.p.* with NaOx (60 mg/kg/day for 3 consecutive days). Kidney stone deposition was examined by Alizarin Red S. Quantification of the affected area in the whole kidney was also shown. Each dot represents the average of the affected areas of 4 whole-mount slides from one mouse. \**P*<0.05, \*\**P*<0.01, \*\*\**P*<0.005, \*\*\*\**P*<0.001 by one-way ANOVA (A) and two-tailed unpaired *t* test (B and C). Data are depicted as mean±SEM. Data are derived from at least 2 independent experiments.

CaPi by itself is also a good nucleator of CaOx(*8*) which is the most prevalent type of kidney stone in human. We next tested whether the excessive presence of urine CaPi in the mutated mice would predispose CaOx stone formation. To this end, we first tested whether high urine Pi burden would accelerate CaOx formation. C57BL/6 mice *i.p.* treated with sodium oxalate (NaOx) for 3 consecutive days resulted in apparent CaOx stone deposition in the kidney at the end of treatment, as revealed by Alizarin Red S staining (Fig. 3B, left). Co-injecting mice with a Pi-rich solution indeed increased CaOx stone formation (Fig. 3B, right). When the tamoxifen-treated *Slc34a1*^ΔMØ^ mice were challenged with the same NaOx regimen, more severe CaOx stone deposition, in particular at the area from the inner cortex to the cortex-medulla junction, was noticed in comparison with vehicle-treated *Slc34a1*^ΔMØ^ mice during NaOx challenge (Fig. 3C). Altogether, these data demonstrate that the expression of SLC34A1 in kidney cortex MØ is crucial in precluding kidney stone formation by controlling the Pi concentration in the urine.

### Macrophages positioned at the proximal tubules of inner cortex are critical to preventing mineral crystal formation by moving Pi from the urine to the kidney interstitium

We noticed that the increased deposition of both CaPi and CaOx crystals in *Slc34a1*^ΔMØ^ mice was particularly prominent in the inner cortex (Figs. 3A-C). This mineral stone distribution pattern was consistent with others’ findings in both CaPi(*35*) and CaOx kidney stone models(*36*), implying that certain microanatomical structure or physiological propensity renders the inner cortex susceptible to the formation of mineral crystals. Proximal tubules constitute the majority of cortical tubules(*37*) and they consist of 3 segments sequentially numbered S1 to S3 with increasing distance from the glomerulus (Fig. 4A)(*38*). S3 segment can be differentiated from S1 and S2 segments by its straight contour and anatomic distribution pattern (spanning the inner cortex and outer medulla)(*39*). High-magnification microscopic examination of the kidneys of *Slc34a1*^ΔMØ^ mice revealed that most CaPi crystals were in the LTL^+^ (Lotus Tetragonolobus Lectin) proximal tubules of inner cortex, particularly at the caudal of S2 segment and the S3 segments (together designated as late proximal tubules hereafter) (Fig. 4B). A major difference of urine between the early and later proximal tubules is a continuous Ca^2+^ concentration rise. Ca^2+^ is predominantly reabsorbed in the proximal tubules via passive paracellular transport(*16*). The early tubules have high sodium and water reabsorption ability but low Ca^2+^ reabsorption ability; active reabsorption of water in the early tubules raises the concentration of urine Ca^2+^ so as to create a concentration gradient that drives paracellular Ca^2+^ reabsorption(*16*) which is further facilitated by a relative high expression of claudin-2, a Ca^2+^ transporter, in the late proximal tubules(*40*). We thus figured that the physiologically high intraluminal Ca²[ concentration made late proximal tubules in the inner cortex susceptible to mineral crystal supersaturation, particularly when there was excessive Pi or oxalate in urine. In this context, MØ-facilitated Pi reabsorption in late proximal tubules could be critical to reduce intraluminal concentration of Pi so as to lower the likelihood of CaPi crystallization.

**Fig. 4.**
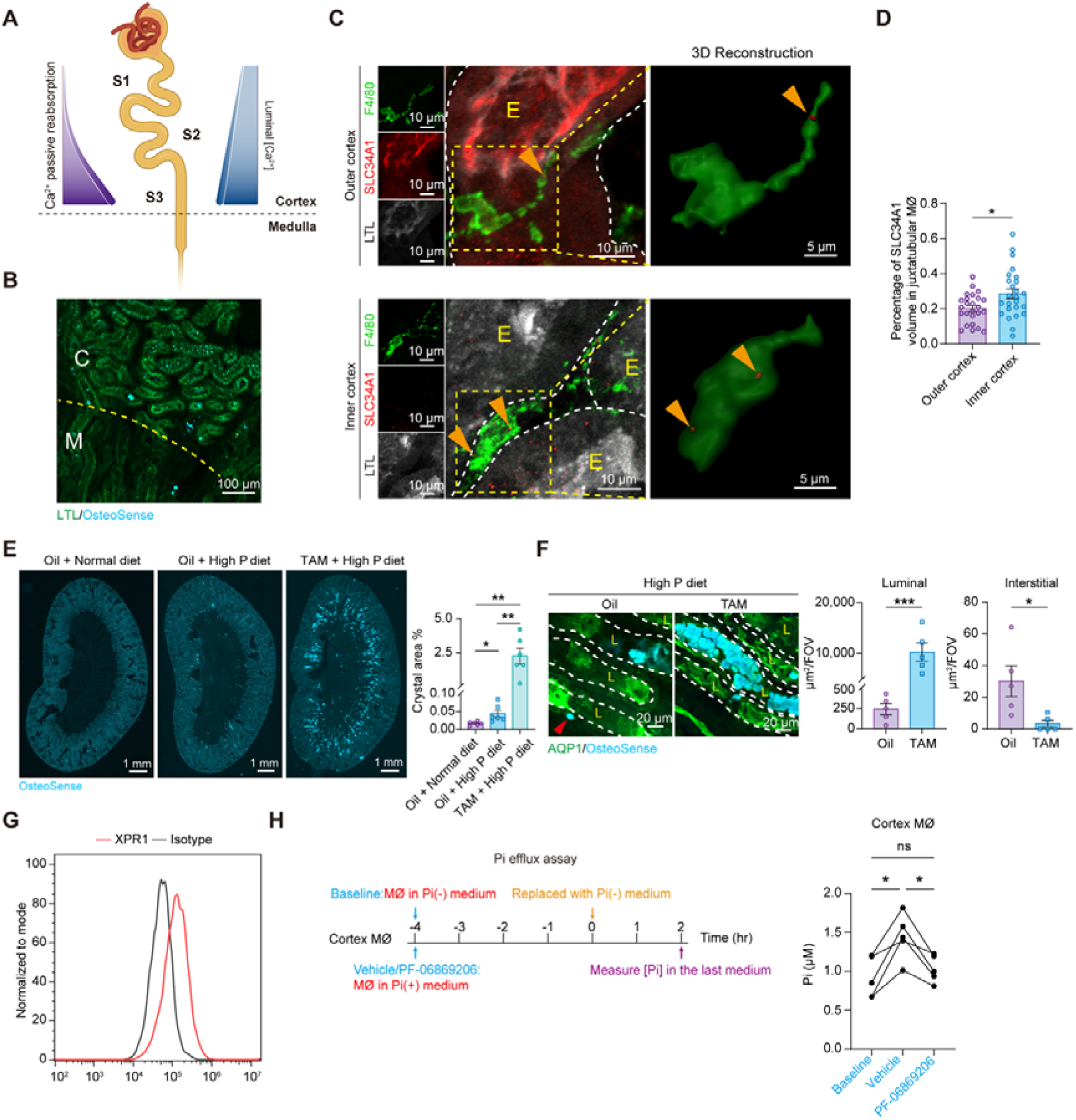
Pi reabsorption by macrophages in the inner cortex is critical to preventing mineral crystal formation. (**A**) Schematic diagram displays that an increased luminal Ca^2+^ concentration in the late proximal tubules drives Ca^2+^ passive reabsorption. (**B**) *Slc34a1*^ΔMØ^ mice (after TAM treatment) displayed CaPi crystal (OsteoSense^+^) deposition particularly in the late proximal tubules the inner cortex. Yellow dotted line: border between cortex and medulla. C, cortex. M, medulla. (**C**) In C57BL/6 mice, SLC34A1 expression in outer and inner cortex was examined by immunohistochemistry. White dotted line, the basolateral border of proximal tubules. E, epithelial cells. Arrowheads, SLC34A1 signal in MØ. (**D**) Statistical comparison of juxtatubular MØ SLC34A1 expression between outer and inner cortex. Each dot indicates the average value of 5 randomly chosen juxtatubular MØ derived from one mouse. n = 5. (**E, F**) *Slc34a1*^ΔMØ^ mice were under normal or high-phosphorus diet for 9 d in the indicated treatment. (**E**) Representative images of OsteoSense staining in the kidneys. The affected areas of CaPi crystal deposition were compared. Each dot represents the average of 4 whole-mount slides from one mouse. (**F**) The positions of CaPi crystals relative to the proximal tubules (AQP1^+^) in the kidney inner cortex of *Slc34a1*^ΔMØ^ mice after the indicated treatment. Dotted line, the border between tubule and interstitium. L, tubule lumen. Arrowhead, CaPi crystal in the interstitium. Each dot indicates the average of CaPi-deposited luminal or interstitial area in four 319 × 319 μm^2^ FOV derived from one mouse. (**G**) A representative histogram of XPR1 expression in cortex MØ. An anti-XPR1 antibody and an isotype antibody was used for flow cytometry analysis. (**H**)(left) Scheme illustrating the protocol of *ex vivo* Pi efflux assay for cortex MØ and (right) statistical analysis of Pi released into the Pi-free medium. PF-06869206, an SLC34A1-specific blocker. *P<0.05, \*\**P*<0.01, \*\*\*\**P*<0.001 by two-tailed unpaired *t* test (D, E, G) and by one-way ANOVA (H). ns, not significant. Data are depicted as mean±SEM. Data are derived from 3 independent experiments.

To interrogate the importance of MØ close to the late proximal tubules in the inner cortex, we first examined the pattern of MØ SLC34A1 expression in the cortex. With an anti-SLC34A1 antibody, immunohistochemical analysis verified a previous reported distribution pattern of SLC34A1 in proximal tubules with an enriched expression in the early convoluted segments but a significantly declined expression in the late straight segment (Fig. 4C)(*41*). Interestingly, confocal analysis of thick sections of the cortex showed that juxtatubular MØ had a converse express pattern to the proximal tubules, as the MØ attached to the late proximal tubules in the inner cortex had a relatively higher SLC34A1 expression than their counterparts adjacent to tubules in the early segments in outer cortex (Figs. 4C and 4D). Thus, MØ with relatively high SLC34A1 expression were distributed in the proximity of late proximal tubules in the inner cortex.

The susceptibility of late proximal tubules to mineral stone formation and the importance of MØ-mediated protection were further highlighted by the mice under a high-phosphorus (2.4%) diet. In the control corn oil-treated *Slc34a1*^ΔMØ^ mice, high-phosphorus diet promoted CaPi supersaturation and crystal deposition, particularly in the inner cortex, by 2.6 folds (Fig. 4E). Pronouncedly, loss of the protection of MØ SLC34A1 in tamoxifen-treated *Slc34a1*^ΔMØ^ mice resulted in a further 50.8-fold increase of CaPi crystal deposition under high-phosphorus diet, when compared to the control mice treated in the same diet (Fig. 4E). In these vulnerable MØ SLC34A1-deprived mice, crystal deposition started remarkably in the proximal tubules of inner cortex and spread to the downstream tubules in the medulla, underscoring an indispensable role of MØ Pi reabsorption in limiting mineral stone formation in the susceptible late proximal tubules.

Tubular epithelial cells balance the reabsorbed urine-derived electrolytes including Pi by releasing them to the kidney interstitium at the basolateral side of the tubules where electrolytes would diffuse to the capillary by concentration gradient. Interestingly, close examination of inner cortex of control mice under high-phosphorus diet revealed that there were CaPi microcrystals present in the interstitium near the late proximal tubules in addition to their intratubular over-presence (Fig. 4F), implying that an over-efflux of Pi excessively reabsorbed by MØ to the interstitium occurred and together with the aforementioned constitutively high efflux of Ca^2+^ from the late proximal tubules, CaPi supersaturated and crystal formed in the interstitium. Supporting this notion that the Pi reabsorbed via SLC34A1 by MØ was released to the interstitium, although high-phosphorus diet induced a 50-fold increase of CaPi crystal deposition in the tamoxifen-treated *Slc34a1*^ΔMØ^ mice (Fig. 4E), the crystals were almost intraluminal without a concomitant increased deposition in interstitium (Fig. 4F). Corroborating this finding, we found that cortex MØ express XPR1, the only known Pi exporter mediating the efflux of Pi in epithelial cells to the interstitium (basolateral side of tubules) (*42, 43*) (Fig. 4G). To test whether cortex MØ did release Pi after reabsorption, we performed a Pi efflux assay by pre-incubating cortex MØ derived from normal mice in normal medium containing Pi for 4 hr. Afterwards, cells were replaced with a Pi-free medium for another 2 hr. We then harvested the last medium and tested the Pi inside which was released by the MØ. MØ were given a SLC34A1 blocker or vehicle in the first 4 hr of Pi-rich incubation. The results clearly showed that cortex MØ would release Pi into the environment, which, however, was substantially suppressed when their Pi uptake ability was pre-blocked, demonstrating that cortex MØ would release the Pi pre-absorbed via SLC34A1 (Fig. 4H).

### MØ SLC34A1 expression is not redundant to tubular epithelial cells but proactively precludes an accelerated mineral stone formation

Our previous results showed that the monocytes would upregulate SLC34A1 expression when infiltrating into the kidney cortex and differentiating into MØ (Fig. 1G). To probe what kidney-associated cue elicited their SLC34A1 expression, we first co-cultured bone marrow-purified monocytes with tubular epithelial cells purified from kidney cortex to examine whether the cues were generated from the tubular epithelial cells which are the primary tissue cells in the kidney. The tubular epithelial cells were verified without cortex MØ contamination before co-culture with monocytes (fig. S4B). However, we did not observe an increased *Slc34a1* expression in monocytes (fig. S8). Previous studies showed that a variety of inorganic particles, including CaPi crystals, could stimulate MØ(*44–46*). The proximal tubules are the primary places of water reabsorption and microcrystals including CaPi particles continuously form in the normal urine(*16*). We thus speculated that CaPi microcrystals could by themselves induce SLC34A1 in monocytes and the differentiated MØ. Indeed, accompanying an over-generation of intratubular CaPi crystals under high-phosphorus diet, upregulation of SLC34A1 expression in the inner cortex MØ was observed in normal mice (Fig. 5A). However, acidifying urine to dissolve CaPi crystals by feeding mice with NH_4_Cl water could abrogate the SLC34A1upregulation in MØ (Fig. 5A). Consistent with this in vivo observation, monocytes responded to CaPi crystals by upregulating *Slc34a1* expression in a crystalline CaPi dose-dependent manner (Fig. 5B). These data together strongly suggest that exposure to CaPi crystals is a trigger of SLC34A1 expression/upregulation in cortex MØ.

**Fig. 5.**
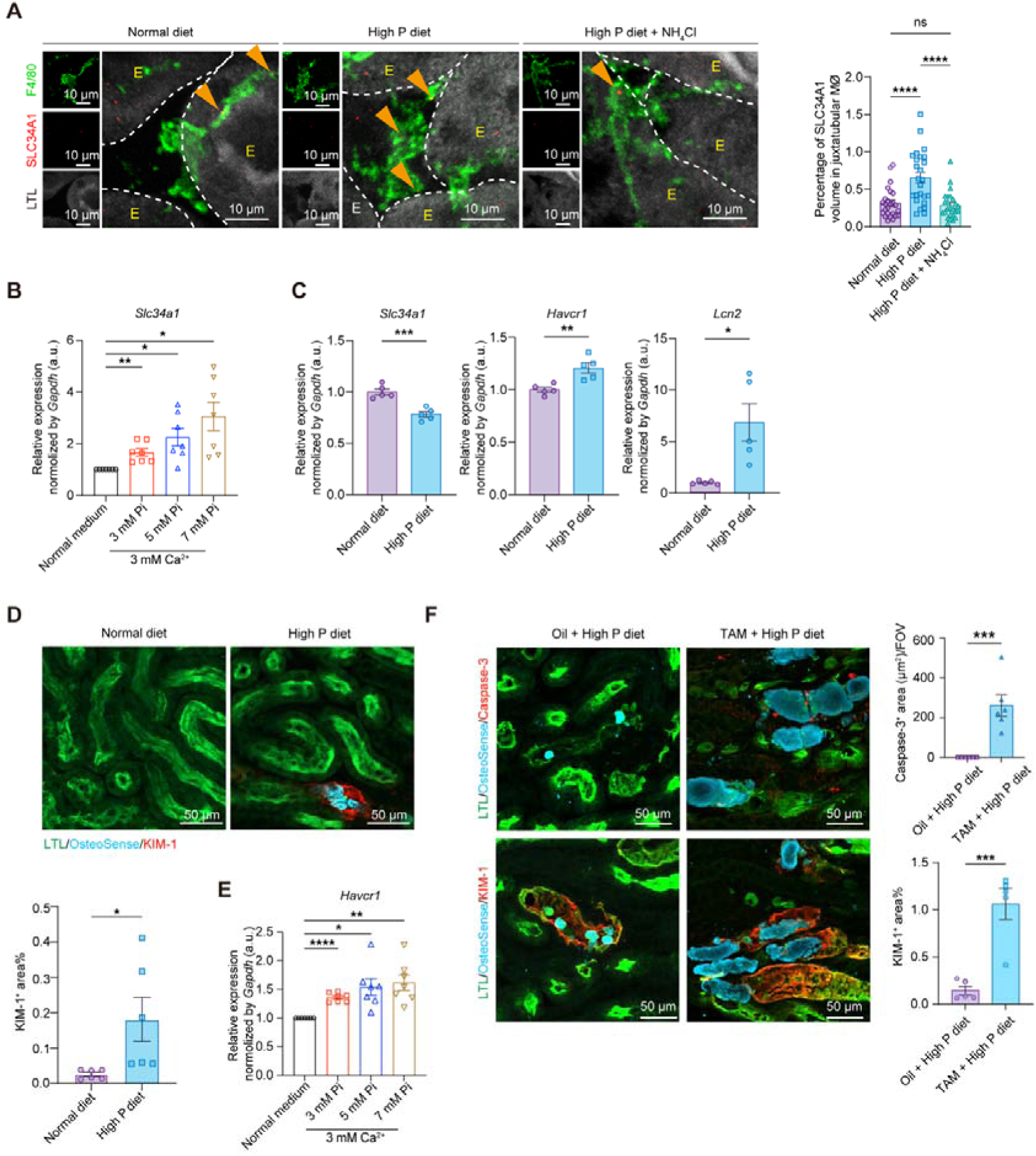
CaPi crystals induce SLC34A1 upregulation in monocytes and kidney cortex MØ but inflict damages to renal tubular epithelial cells. (**A**) C57BL/6 mice were treated with high-phosphorus diet for 9 d, and SLC34A1 expression in the juxtatubular MØ of inner cortex was compared. Each dot indicates the average value of 5 randomly chosen juxtatubular MØ derived from one mouse. n = 5. (**B**) Bone marrow-purified monocytes were cultured in normal medium or medium supplemented with Pi and Ca^2+^ to the indicated amounts for 6 hr. Expression of *Slc34a1* was measured by RT-PCR. (**C**) The transcript expression of *Slc34a1*, *Havcr1* and *Lcn2* in the purified cortex tubules from C57BL/6 mice under normal or high-phosphorus diet. (**D**) Representative images of kidney cortex of C57BL/6 mice under normal or high-phosphorus diet, which show KIM-1 expression in the proximal tubule epithelial cells (LTL^+^) around CaPi crystals (OsteoSense^+^) under high-phosphorus diet. Statistics shows the KIM-1^+^ area. Each dot represents the average of the affected areas of two whole-mount slides from one mouse. (**E**) Purified cortical tubular cells were cultured in normal medium or medium supplemented with Pi and Ca^2+^ to the indicated amounts for 24 hr. Their relative expression of *Havcr1* was evaluated by RT-PCR. (**F**) Representative images and statistics of caspase-3 staining (upper panels) and KIM-1 staining (lower panels) showing degrees of tubular epithelial injury under the indicated treatment of *Slc34a1*^ΔMØ^ mice. Caspase-3: Each dot represents the average of 4 randomly chosen FOV in the size of 319 × 319 μm^2^ derived from one mouse. n = 6. KIM-1: Each dot represents the average of the affected areas of two whole-mount slides from one mouse. n = 5. \**P*<0.05, \*\**P*<0.01, \*\*\**P*<0.005, \*\*\*\**P*<0.001 by one-way ANOVA (B and E) and two-tailed unpaired *t* test (C and D). Data are depicted as mean±SEM. Data are derived from at least 2 independent experiments.

In contrast, cortex tubular epithelial cells derived from wild-type mice under high-phosphorus diet displayed a downregulated *Slc34a1* expression, accompanied with an increased expression of renal epithelial injury markers *Havcr1* and *Lcn2* which encode KIM-1 and NGAL, respectively(*47–49*) (Fig. 5C). Histologically, we could conspicuously detect KIM-1^+^ injured tubular epithelial cells around the CaPi crystals in mice intaking high-phosphorus diet (Fig. 5D). *In vitro*, CaPi microcrystals would induce *Havcr1* expression in primary cortical tubular epithelial cells in a dose-dependent manner (Fig. 5E). We previously showed that when MØ *Slc34a1*-deprived mice were under a high-phosphorus diet, a drastic stone formation was noticed (Fig. 4G). This was accompanied by even more severe epithelial injury, manifested by tubule cell apoptosis, which was not observed when MØ *Slc34a1* expression was intact (Fig. 5F, upper panels). Unsurprisingly, KIM-1 expression was more widespread in MØ *Slc34a1*-deprived mice compared to *Slc34a1*-intact mice under high-phosphorus diet (Fig. 5F, lower panels). These data demonstrate that kidney tubule injury was positively correlated with the degree of mineral stone generation; in a situation with an increased intratubular CaPi crystals, cortex MØ would respond by upregulating SLC34A1 expression to enhance Pi reabsorption so as to limit further CaPi supersaturation. Without this proactive response of cortex MØ, renal tissue integrity would be more vulnerable to be breached. Thus, the expression of SLC34A1 by cortex MØ is not redundant to tubular epithelial cells in two-fold aspects: (1) it could compensate a reduced Pi reabsorption ability when tubular epithelial cells were injured by kidney stone formation; and (2) it also proactively precludes an accelerated mineral stone formation.

### Lysosomal damage mediates SLC34A1 upregulation in macrophages

Lysosome is the primary organelle responsible for the degradation of extracellular materials taken up by MØ. We thus speculated that the crystal-related geometric shape of CaPi might impair the lysosomes upon phagocytosis, which may generate signal(s) for *Slc34a1* upregulation. To interrogate lysosomal integrity, we first measured LysoTracker signal intensity by flow cytometry (*50, 51*). We found that treatment of kidney cortex MØ *ex vivo* with prepared CaPi particles markedly reduced LysoTracker signal intensity (Fig. 6A), suggesting lysosome impairment. Consistently, we observed a decrease in total LAMP1^+^ lysosome volume in cortex juxtatubular MØ on per cell basis under a high-phosphorus diet (Fig. 6B), a treatment which would increase CaPi deposition in the kidney. Since monocytes were derived from bone marrow in adults, we next employed bone marrow culture-derived MØ (BMDMs) to explore the mechanisms of CaPi-induced *Slc34a1* expression. As expected, we could recapitulate an upregulation of *Slc34a1* expression and a decrease in lysosomal integrity upon CaPi particle treatment in BMDMs (Fig. 6C, D). Concurrently, CaPi treatment raised lysosomal pH, indicated by LysoSensor(*52*) staining (Fig. 6E), another index of lysosome dysfunction.

**Fig. 6.**
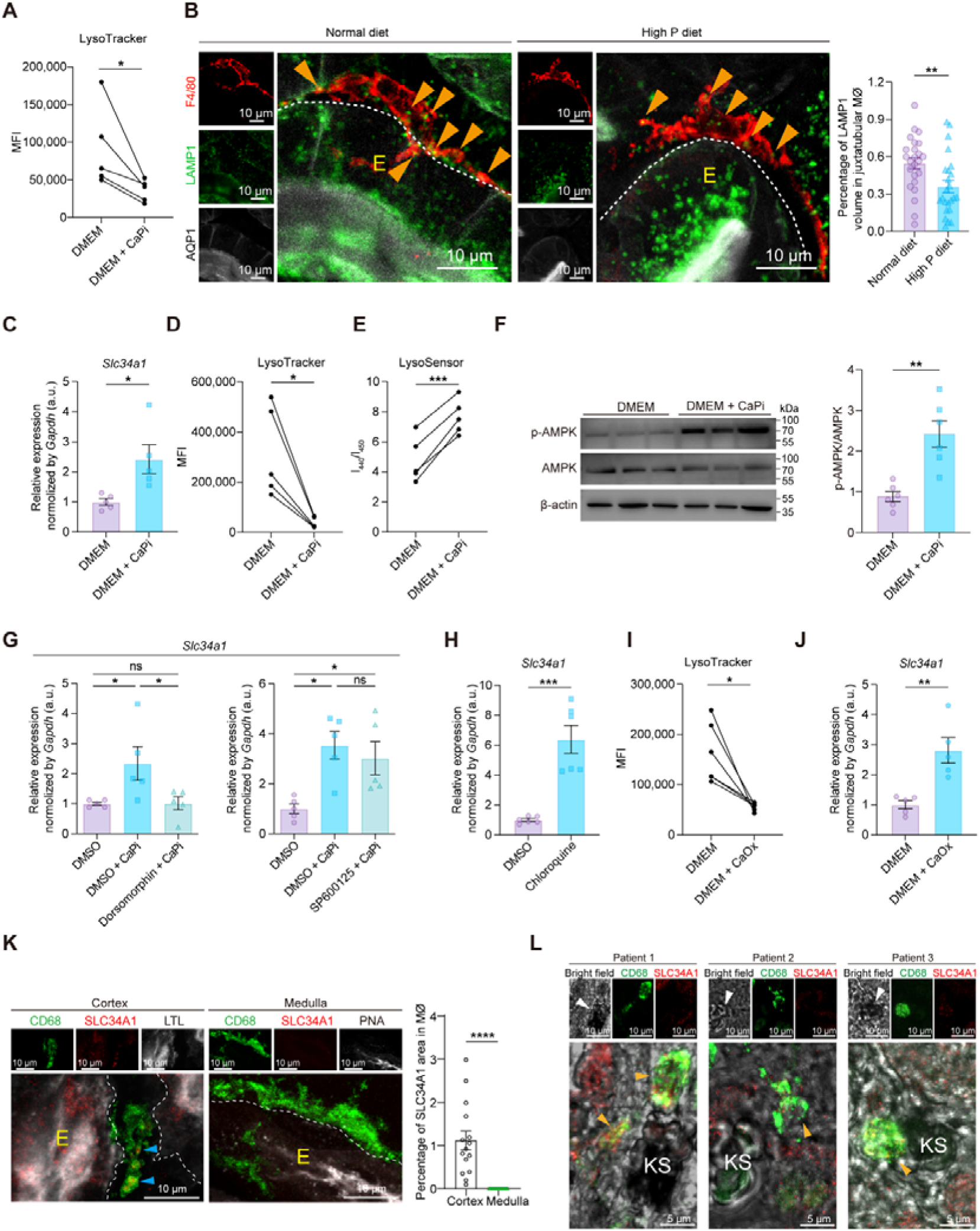
Lysosome damage by mineral microcrystals induces SLC34A1 upregulation in macrophages. (**A**) Kidney cortex MØ were treated *ex vivo* with or without prepared CaPi particles for 2 hr and then stained with LysoTracker. MFI of LysoTracker was examined by flow cytometry analysis. (**B**) Representative images of kidney cortex of C57BL/6 mice under normal or high-phosphorus diet for 6 d, which show LAMP1 expression in the juxtatubular MØ. Each dot indicates the average value of MØ from one 319 × 319 μm^2^ FOV. These images also exhibit that epithelial cells highly expressed LAMP1, as reported before(*66*). Arrowhead, LAMP1 signal in MØ. Dotted line: the basolateral border of tubule. E, epithelial cells. n = 6. (**C**-**E**) BMDMs were treated with or without prepared CaPi particles for 2 hr. (**C**) Expression of *Slc34a1* was measured by RT-PCR. (**D**) Cells were stained with LysoTracker and the intensity of LysoTracker was examined. (**E**) Cells were stained with LysoSensor and the intensity of LysoSensor was examined. (**F**) BMDMs were treated with or without prepared CaPi particles for 30 min. The abundance of phosphorylated AMPK (p-AMPK) and AMPK was measured by Western blot and the statistics show their ratio. (**G**) BMDMs were pre-treated with AMPK inhibitor dorsomorphin (5 μM) for 30 min or JNK inhibitor SP600125 (20 μM) for 1 hr, followed by being cultured with or without prepared CaPi particles for 2 hr. Some cells were pre-treated with vehicle (DMSO). Expression of *Slc34a1* was then measured by RT-PCR. (**H**) BMDMs were treated with or without chloroquine (100 μM) for 2 hr. Expression of *Slc34a1* was then measured by RT-PCR. (**I** and **J**) BMDMs were treated with or without prepared CaOx particles for 2 hr. (**I**) Cells were stained with LysoTracker and the intensity of LysoTracker was examined. (**J**) Expression of *Slc34a1* was measured by RT-PCR. (**K**) Representative images and statistical analysis of SLC34A1 expression in cortex and medulla MØ of human kidney paracancerous tissues. Each dot represents the relative SLC34A1 expression in one MØ. n=3. LTL labels cortex proximal tubules. PNA labels medulla collecting ducts. Arrowhead, SLC34A1 signal in MØ. (**L**) Representative images derived from 3 patients of nephrolithiasis showed that cortex MØ which surrounded kidney stones highly express SLC34A1. White arrowhead, position of kidney stone. Yellow arrowhead, SLC34A1-expressing MØ. KS, kidney stone. n.s. not significant. \**P*<0.05, \*\**P*<0.01, \*\*\**P*<0.005, \*\*\*\**P*<0.001 by two-tailed paired *t* test (A, D, E and I), two-tailed unpaired *t* test (B, C, F, H, J and K), and one-way ANOVA (G). Data are depicted as mean±SEM. Data are derived from at least 2 independent experiments.

Lysosome is known for functioning as a homeostasis-regulating hub and its integrity disruption will activate AMP-activated kinase (AMPK) signaling(*53*). In accord, we found that stimulation with CaPi crystals led to an increased p-AMPK/AMPK ratio in MØ (Fig. 6F). Pre-treatment with dorsomorphin, a potent AMPK inhibitor, could abrogate MØ of *Slc34a1* upregulation upon exposure to CaPi crystals (Fig. 6G, left panel). A recent report showed that particle uptake by MØ could activate C-Jun N-terminal kinases (JNKs) in addition to AMPK(*46*). However, pre-treating MØ with a JNK inhibitor did not alter the effect of CaPi crystals on *Slc34a1* upregulation (Fig. 6G, right panel). These data suggest that it was lysosome disruption-mediated AMPK activation that induced *Slc34a1* upregulation upon MØ encounter with CaPi crystals. Corroborating this notion, pharmacological treatment of MØ with chloroquine, an agent well known for interfering with lysosome function and stimulating AMPK(*54, 55*), could also lead to *Slc34a1* upregulation in MØ (Fig. 6H).

Besides CaPi, CaOx manifests as another common type of mineral crystals present in renal tubules. Interestingly, we observed that like CaPi, CaOx treatment could also impair lysosome integrity and increase *Slc34a1* expression in MØ (Figs. 6I and 6J). Thus, the two most prevalent nephrolithiasis-relevant crystals could both stimulate an upregulation of SLC34A1 in MØ. Together with the data that mice without MØ *Slc34a1* expression were more vulnerable to develop CaOx stones (Fig. 3C). SLC34A1 expression by cortex MØ could be considered as a general strategy to reduce mineral crystal formation in the downstream renal tubules.

In human, renopuncture in most cases only samples the cortex but not the medulla. To assess whether MØ in human kidney also have disparity in SLC34A1 expression between cortex and medulla, we analyzed paracancerous kidney tissues removed from surgery, which provided both cortex and medulla in a relative normal state. It revealed that similar to that in mice, juxtatubular MØ in the cortex but not those in the medulla express SLC34A1 (Fig. 6K). In renopuncture samples derived from patients with nephrolith, MØ were noted to surround mineral crystals and they highly express SLC34A1 (Fig. 6L).

## Discussion

During the filtrate of the glomerulus flows though the renal tubular system, mineral crystals resulting from urine concentration generate numerous microscopic sedimental particles, which places an imminent threat to tubule clearance and nephron function. CaOx and CaPi stones are the two most prevalent kidney stones, and they are frequently formed as mixtures(*7, 8*). Thus, it is conceivable that limiting the concentrations of oxalate, Ca^2+^ and/or Pi in the urine could be a practicable way to reduce mineral stone formation. In the kidney, there is no reabsorption transporter for oxalate which has an absolute excretion by the urine(*11*); in term of Ca^2+^, the majority of Ca^2+^ is reabsorbed through a passive paracellular pathway relying on the transepithelial electrochemical gradient and ion permeability through tight junctions between renal tubular epithelial cells(*16, 56*). Thus, employing oxalate or Ca^2+^ reabsorption as a strategy to prevent supersaturation of urinary minerals is limited. In the current study, we found that kidney cortex MØ distinguishably express SLC34A1 and could reabsorb Pi from the urine as a means to reduce the risk of CaPi crystal formation in the tubules. Since CaPi also serves nucleator for CaOx growth, reducing the amount of Pi in the urine also limits CaOx crystal formation. The preventive strategy employed by MØ is proactive since they upregulate SLC34A1 expression on encountering mineral crystals in the urine. Deprivation of *Slc34a1* expression in kidney cortex MØ led to not only an over-presence of Pi and Ca^2+^ in the urine, but a markedly expedited CaPi microparticle deposition in kidney tubules as well. As such, an over-supply of NaOx on top of that resulted in an accelerated kidney stone formation, substantiating an indispensable role of kidney MØ in preventing a supersaturation of CaPi in the urine and the formation of kidney stones.

We pinpointed in this study that the late proximal tubules in the inner cortex are vulnerable to mineral stone formation. Based on the current study and our previous finding, there are two-fold reasons: (1) the physiological high intraluminal concentration of Ca^2+^ predisposes to supersaturation of mineral crystals if an excessive Pi or oxalate is present in the urine; (2) unlike medulla MØ, cortex MØ are ill-equipped with capability in transmigration and cleaning intratubular large crystals(*20*). To compensate to the above weakness, the MØ adjacent to these inner cortex tubules are highly capable in reabsorbing Pi via SLC34A1 expression, another strategy to prevent mineral stone formation. In addition, this finding is the first report of a non-epithelial cells in reabsorbing electrolytes in the urine. The importance of MØ but not tubular epithelial cells in modulating Pi concentration in the urine was underscored by their different responses to CaPi microcrystals. Over-presence of mineral crystals in the tubules would injure tubular epithelial cells, as shown by us in the current study and others(*34*), which led to a downregulation of SLC34A1 expression; in contrast, MØ would be stimulated upon encountering mineral crystals(*44–46*) and upregulate SLC34A1 expression as a response. Thus, in addition to a previous finding about a role of medulla MØ in the clearance of preformed intratubular particles(*20*), here we identified that the upstream cortex MØ opted to enact another strategy of deaccelerating mineral particle deposition by actively limiting Pi concentration in the urine. These newly defined behaviors of kidney MØ highlight their physiological roles as the previously unappreciated guardians to maintain renal tubule unblocked. Current treatments for kidneys stones are limited to surgery/scope removal, ultrasound breaking-up, and enhancing urine flushing or stone dissolving by increasing daily water intake volume and adjusting drinking water pH. The cellular mechanisms involving MØ should be explored in the future for therapeutical application on kidney stone diseases.

MØ functions are imprinted by multiple factors, and increasing evidence suggests that microenvironment plays a critical role(*57, 58*). The surrounding tissue cells construct distinctive niches for tissue-specific resident MØ by providing not only growth factors, such as CSF1(*59*), but instructing factors to guide MØ differentiation and activation which in turn execute tailored functions to suffice the needs of surrounding cells(*60*). This reciprocal relationship between MØ and their resided milieu maintain tissue homeostasis and functions.

Kidney medulla MØ are embryo-derived and they interact with collecting ducts *via* integrin-fibronectin interaction(*20*). In the case of kidney cortex MØ, bone marrow-derived monocytes routinely infiltrate to the cortex and differentiate to MØ(*20*). Our adoptive transfer experiment showed that monocytes upregulated SLC34A1 in the kidney but not in the liver or spleen when infiltrating into tissues and differentiating into MØ, implying that kidney cortex-specific local milieu is accountable for this change. However, mineral crystals also exist in kidney medulla but we show here in both transcriptomic study and SLC34A1 reporter mice that medulla MØ do not express SLC34A1. Whether it is due to its embryo-origin or other microenvironment-imprinting effects is unknown yet. It is well appreciated that the microenvironments between the cortex and medulla are very different in that there distribute different tubular segments with distinctive functions and the medulla is relatively hyperosmotic and hypoxic(*61–63*). Whether these milieu cues also drive the morphological and functional characteristics of macrophages in the medulla deserves further exploration.

## ACKNOWLEDGMENTS

We thank Jingyao Chen from the Core Facilities, Zhejiang University School of Medicine for the technical support. We also thank Qin Han for her technical assistance on Confocal Microscopy in the Center of Cryo-Electron Microscopy (CCEM), Zhejiang University.

## AUTHOR CONTRIBUTIONS

Conceptualization, X.Z.S.; designing, F.H. and X.Z.S.; supervision, F.H. and X.Z.S.; imaging, Yuxi W., Yuancheng W. and X.D.; RNA-seq and bio-informatics analyses, Yuancheng W.; blood biochemical analysis, Q.W.; human sample preparation, F.X. and D.L.; all other experiments, Yuxi W., Yuancheng W., X.D., N.C., Z.L., Z.G. and J.H.; statistics, Yuxi W. and Yuancheng W.; validation, F.H.; writing, X.Z.S.

## DATA AVAILABILITY

Data will be made available on request. The RNA-seq data of renal cortex and medulla macrophages reported in this study have been deposited in the Genome Sequence Archive database (https://ngdc.cncb.ac.cn/gsa/), and the accession number is GSA: CRA007988.

## MATERIALS AND METHODS

### Study approval

All animal experiments adhered to the NIH Guide for the Care and Use of Laboratory Animals, and were approved by the Institutional Animal Care and Use Committee at Zhejiang University (ZJU20240398). Processing of human samples and data was approved by the Jinling Hospital Ethics Committee (2025DZKY-072-01), and informed consent was obtained from all participants and legal guardians. The study design and conduct complied with all relevant regulations regarding the use of human research participants and was conducted in accordance with the criteria set by the Declaration of Helsinki.

### Animals

C57BL/6 mice and *Slc34a1*^CreERT2/+^ mice (NM-KI-210110) were purchased from Shanghai Model Organisms Center, Inc.; *Cx3cr1*^CreERT2/+^ mice (020940), Ai14 (*Rosa26-stop-TdTomato*) mice (007914) and *Rosa26-*iDTR mice (007900) were obtained from The Jackson Laboratory; *Slc34a1*^fl/fl^ (T019090) and CD45.1 (T054816) mice, both of which are at C57BL/6 background, were originally purchased from GemPharmatech. Mice were housed in a standard animal facility, with a 12-hr light/dark cycle, in specific-pathogen-free environment. Our study examined male and female animals, and similar findings are reported for both sexes.

To induce deletion of targeted genes, adult mice received *i.p.* injection of 60 mg/kg (body weight) tamoxifen (MedChemExpress, #S7818) daily for 5 consecutive days or regimens specified in experimental protocols. For mouse kidney capsule injection, the procedure typically involves anesthetizing the mouse and making an incision to expose each kidney. Utilizing a 30-gauge syringe needle, carefully lift the kidney capsule and inject 40 μl of 30 mg/ml tamoxifen per kidney.

For low-phosphorus and high-phosphorus diet treatment, mice were fed with a diet containing 0.02% and 2.4% inorganic phosphorus, respectively.

To acidify the urine, add 3% ammonium chloride (NH_4_Cl) to the drinking water of mice for 1 week.

PF-06869206 (MedChemExpress, #HY-112065), an SLC34A1 specific inhibitor, was administered *i.p.* at a dose of 300 mg/kg.

### Public transcriptomics data analysis

We analyzed two publicly available RNA-seq datasets. The first was the kidney cortex and medulla macrophage bulk RNA-seq data previously reported by us(*20*). We used R package DESeq2 (v1.34.0) to do differential gene expression analysis(*64*), during which process we excluded genes with less than 30 total mapped reads among the 6 samples, and kept genes that had more than 10 mapped reads in no less than 2 samples in either group. Finally, we got 1,517 upregulated genes and (log2FC > 1, adjusted *P* value < 0.001) and 632 downregulated genes (log2FC < -1, adjusted *P* value < 0.001) in the cortical macrophages vs. medullary macrophages. Then we performed GO analysis with the DEGs *via* package clusterProfiler (v4.0.5)(*65*). The *enrichGO* function was used with parameters: ‘OrgDb’ = ‘org.Mm.eg.db’, ‘ont’ = ‘BP’, ‘pvalueCutoff’ = 0.05, ‘qvalueCutoff’ = 0.05, and ‘pAdjust-Method’ = ‘BH’ (Benjamin and Hochberg correction). Then *simplify* function was used to remove redundant GO terms with parameters: cutoff = 0.7, by = “p.adjust”, select_fun = min. For analyzing renal macrophages gene expression ranking, we first obtained the gene expression values (TPM) after quantification. After filtering out the genes that had no expression (TPM = 0 in all the samples), we ranked the remaining genes according to their TPM values in individual samples or their average TPM of the three samples in each group. The second was RNA-seq data about resident macrophages derived from multiple organs, which was collected by Friedrich et al.(*23*). We obtained the normalized data using the robust multi-array average by the authors; phosphate transporter genes expression levels in macrophage were determined by averaging the normalized gene expression.

### Real-time PCR

Renal cortex macrophage and proximal tubular fragments were lysed separately using Trizol reagent, followed by the addition of chloroform (Sinopharm Chemical Reagent Co.,Ltd, #10006818) and centrifugation. The aqueous phase was collected, and RNA was precipitated with isopropanol (Sinopharm Chemical Reagent Co.,Ltd, #80109218). The RNA pellet was then washed with 75% ethanol (Sinopharm Chemical Reagent Co.,Ltd, #801769610) and finally dissolved in DNase/RNase-Free water (Coolaber, #SL2250). And RNA extraction from tissues was performed using a kit from ES Science (#ES-RN002plus). cDNA synthesis was performed using 1 μg of purified RNA with a high-Capacity cDNA reverse transcription kit (Yeasen, #11141ES60). The mRNA levels of each target were analyzed by quantitative real-time PCR using SYBR Green Mix (Yeasen, #11201ES08) and the CFX96 Touch Real-Time PCR Detection System (Bio-Rad). Gene expression levels were normalized to the internal control gene *Gapdh*. The following primers were used: *Gapdh* forward primer AGGTCGGTGTGAACGGATTTG *Gapdh* reverse primer TGTAGACCATGTAGTTGAGGTCA *Aqp1* forward primer AGGCTTCAATTACCCACTGGA *Aqp1* reverse primer GTGAGCACCGCTGATGTGA *Slc9a3* forward primer TGAAAAGCAGGACAAGGAAATCT *Slc9a3* reverse primer TTGGCCGCCTTCTTATTCTGG *Cyp27b1* forward primer AGAGCGCTGTAGTTTCTCATCATG *Cyp27b1* reverse primer CCATCCGCCGTTAGCAAT *Cyp24a1* forward primer ACCCCCAAGGTCCGTGACATC *Cyp24a1* reverse primer CCAGTTGGTGGGTCCAGGTAAGG *Tnf*α forward primer AAGCCTGTAGCCCACGTCGTA *Tnf*α reverse primer GGCACCACTAGTTGGTTGTCTTTG *Il1*β forward primer ACCTTCCAGGATGAGGACATGA *Il1*β reverse primer CTAATGGGAACGTCACACACCA *Il6* forward primer TAGTCCTTCCTACCCCAATTTCC *Il6* reverse primer TTGGTCCTTAGCCACTCCTTC *Havcr1* forward primer CTGGAATGGCACTGTGACATCC *Havcr1* reverse primer GCAGATGCCAACATAGAAGCCC *Lcn2* forward primer GAAATATGCACAGGTATCCTC *Lcn2* reverse primer GTAATTTTGAAGTATTGCTTGTTT *Slc34a1* forward primer TGACCCACTACCTACCAAGC *Slc34a1* reverse primer AGGTAAAGGAAAGCCAGCATCA

### Immunohistochemistry

Mice were deeply anesthetized by isoflurane and perfused transcardially with cold PBS, followed by 4% paraformaldehyde (PFA). Kidneys were then removed, fixed overnight in 4% PFA, and then dehydrated in PBS containing 30% sucrose (Sinopharm Chemical Reagent Co.,Ltd, #10021418) at 4 °C for 24 hr. Coronary sections were sliced into 30 μm using a cryostat (Leica, SM2010R). After kidney slices were blocked with the blocking buffer (5% donkey serum (Solarbio, #SL050) and 1% Triton X-100 (Vetec, #V900502-100ML) in PBS) for 2 hr at room temperature, slices were incubated with primary antibody in antibody dilution buffer (0.5% donkey serum and 0.1% Triton X-100 in PBS) at 4 [ overnight. For detection, slices were incubated with secondary antibodies at room temperature in dark for 2 hr. After staining, sections were mounted on slides with Fluoromount-G anti-fade medium containing DAPI (Southern Biotech, #0100-20). The following antibodies were used: rabbit anti-mouse CoraLite^®^ Plus 488-conjugated AQP1 (4B2E10) (Proteintech, #CL488-20333); rabbit anti-mouse SLC34A1 (Novus Biologicals, #NBP2-13328); rat anti-mouse F4/80 (BM8) (ThermoFisher Scientific, #14-4801-81); ABflo^®^ 488 Rabbit anti-Mouse LAMP1 (ABclonal, #A24362); Lotus Tetragonolobus Lectin (LTL)-Fluorescein (Vector Laboratories, #FL-1321-2); goat anti-mouse KIM-1 (R&D Systems, #AF1817); Rabbit anti-Mouse Cleaved Caspase-3 (5A1E) (Cell Signaling Technology, #9664); donkey anti-rabbit AF568 (Abcam, #ab175470); donkey anti-rat AF488 (Abcam, #ab150153); donkey anti-rat AF568 (Abcam, #ab175475); donkey anti-rat AF647 (Abcam, #ab150155); donkey anti-goat AF568 (ThermoFisher Scientific, #A11057).

Staining of human kidney tissue samples was performed on paraffin sections mounted on glass slides. Sections were incubated at 60°C for 1 h, deparaffinized in xylene, and rehydrated through graded ethanol solutions (100%, 90%, and 70%). Heat-induced epitope retrieval was performed in pH 6.0 retrieval buffer. Sections were incubated overnight at 4°C with primary antibodies against CD68 (Santa Cruz, #sc-20060), SLC34A1 or biotinylated LTL (Vector Laboratories, #B-1325-2) and PNA (Vector Laboratories, #B-1075-5), followed by incubation with appropriate secondary antibodies or streptavidin-conjugated fluorophore.

### Confocal microscopy

Confocal z-stacks were captured using LSM800 confocal microscope (Carl Zeiss) equipped with 10×/0.45 NA; 20×/0.8 NA; 40×/0.95 NA and 63×/1.4 NA objective lens or STELLARIS 5 confocal microscope (Leica Microsystems) equipped with 10×/0.40 NA; 20×/0.75 NA; 40×/0.95 NA and 63×/1.4 NA objective lens. Images were analyzed with Imaris software (Bitplane) and Fiji software (NIH).

### Tissue disintegration and flow cytometry analysis

Mice were perfused with 4 [ PBS. Kidney cortex, liver, spleen and brain were separately harvested and first mechanically dissociated in RPMI 1640 medium (HAKATA, #A19006) on ice, followed by enzymatic digestion (Collagenase IV (Worthington, #LS004189) (1.5 mg/ml for kidney, liver and spleen; 0.6 mg/ml for brain) plus 100 U/ml DNase I (Sigma-Aldrich, #DN25) in RPMI 1640) at 37 [ for 30∼60 min. Enzymatic reaction ceased with supplementation of equal volume of 4 [ RPMI 1640 and the digested tissues were filtered through a 70-μm cell strainer (Biosharp, #BS-70-CS). After centrifugation, kidney, liver and brain cell pellets underwent density gradient centrifugation with 36%/72%, 40%/70% and 37%/70% Percoll^TM^ (Cytiva, #17089109) gradients, respectively. Immune cells were enriched at the interfaces of Percoll gradients and were collected. Cells were then subject to flow cytometry (Agilent, Novocyte; Beckman Coulter, CytoFlex S) analysis. Data were processed with FlowJo (v10).

The following antibodies were used for flow cytometry analysis: rabbit anti-mouse CoraLite_®_ Plus 488-conjugated AQP1, PE/Cyanine7 anti-mouse CD115 (AFS98) (Biolegend, #135523), APC anti-mouse/human CD11b (M1/70) (Biolegend, #101211), Pacific Blue anti-mouse/human CD11b (M1/70) (Biolegend, #101223), PE anti-mouse/human CD11b (M1/70) (Biolegend, #101207), BV510 anti-mouse CD45 (30-F11) (Biolegend, #103137), FITC anti-mouse CD45 (30-F11) (Biolegend, #103107), APC anti-mouse CD45.1 (A20) (Biolegend, #110713), Pacific Blue anti-mouse CD45.2 (104) (Biolegend, #109819), BV510 anti-mouse CX3CR1 (SA011F11) (Biolegend, #149025), APC/Fire750 anti-mouse F4/80 (BM8) (Biolegend, #123152), Pacific Blue anti-mouse Ly-6C (HK1.4) (Biolegend, #128013). XPR1 Rabbit PolyAb (Proteintech, #14174-1-AP) and Rabbit IgG control polyclonal antibody (Proteintech, #30000-0-AP); Donkey Anti-Rabbit AF647 (Abcam, #ab150075) was used for secondary staining.

### Kidney cortex macrophage purification

Kidney cortex was separated from the medulla under anatomic microscopy. After aforementioned tissue disintegration, immune cells were first enriched with Percoll gradient centrifugation (as detailed above). The cells were then resuspended in MACS buffer (D-PBS plus with 2%FBS and 1mM EDTA). Following incubation with an Fc blocker (Biolegend, #156603), the cells were incubated with an APC-conjugated anti-mouse F4/80 (BM8) antibody (BioLegend, #123115) on ice for 15 min and then washed with MACS buffer. Subsequently, anti-APC Nanobeads (BioLegend, #488072) were added, and the cells were incubated for an additional 15 min on ice. Afterwards, macrophages were positively collected with a magnet and washed with PBS.

### Monocytes adoptive transfer and fate mapping

Monocytes were isolated from the bone marrow of *Slc34a1*^CreERT2/+^:*R26*^tdTomato^ mice (CD45.2 background) using a monocyte isolation kit (Stemcell Technologies, #19861) based on a negative selection strategy. 3×10^6^ monocytes were then adoptively transferred *i.v.* into each CD45.1/CD45.2 recipient mouse. Starting on the 2^nd^ d post-transfer, recipients were administered tamoxifen daily for seven consecutive days. On the 9^th^ day post-transfer, kidneys, livers, and spleens were harvested and tdTomato expression in macrophages was analyzed by flow cytometry.

### ^32^P uptake Assay

Purified renal cortex macrophages were seeded into a 24-well plate. After a 1.5 hr-incubation at 37 °C with a full medium (RPMI 1640 supplemented with 10% FBS, 1% penicillin/streptomycin), macrophages firmly attached to the plate. Next, 0.4 μM PF-06869206 or DMSO was added into the wells and incubate for another hour. The wells were washed twice with phosphate-free cell culture medium to remove any residual inhibitor or DMSO. Then, 500 μl of 100 μM Na_2_HPO_4_/NaH_2_PO_4_ buffer (containing ^32^P (Revvity, #NEX053001MC) at 1 μCi/ml, prepared in phosphate-free medium (Gibco, #11971025)) was added into each well and incubated for 15 min. Afterwards, the wells were washed by four times with phosphate-free cell culture buffer to remove unincorporated ^32^P, ensuring no radioactive signal remained in the wash solution. The cells were then treated with Trizol reagent (TaKaRa, #9109) and the lysates were transferred into scintillation vials. 15 ml of scintillation cocktail (7 g/L 2,5-Diphenyloxazole, 0.5 g/L 1,4-Bis(5-phenyl-2-oxazolyl) benzene, 650 ml/L Xylene, 350 ml/L Ethylene glycol monomethyl ether) was added into each vial and the ^32^P radioactivity was measured using a liquid scintillation counter.

### Pi efflux assay

Purified renal cortex MØ were first pre-incubated in a 12-well plate with Pi-free DMEM (Gibco, #11971025, for baseline assessment) or normal DMEM (Gibco, #C11995500BT) which contains Pi in the presence of 0.4 μM PF-06869206 or vehicle (DMSO) for 4 hr. Afterwards, medium was replaced with fresh Pi-free medium and the medium was harvested after 2-hr incubation. The concentration of Pi in the medium was determined by malachite green phosphate detection kit (Beyotime, #S0196S). Each sample was measured in duplicate.

### Blood and urine biochemical analyses

Blood was collected *via* retro-orbital sampling, and plasma was isolated by centrifuging at 1,500 g for 15 minutes at 4 °C. Urine samples were collected from mice housed in metabolic cages for the time as indicated. The collected urine was always kept on ice. The urine samples were centrifuged at 1,000 g for 20 min at 4 °C, and the supernatants were collected.

The concentrations of Pi, Ca²[, Na[, K[, Cl^-^ and creatinine in blood plasma and urine were measured by Cobas 8000 ISE and Cobas c702.

### PTH and FGF23 measurements

Plasma levels of PTH and FGF23 were measured by ELISA kits (Elabscience, #E-EL-M0709; Elabscience, #E-EL-M2415, respectively), following the manufacturer’s instructions.

### Measurement of glomerular filtration rate (GFR)

GFR was assessed in conscious, unrestrained mice by monitoring the excretion kinetics of a single intravenous bolus of FITC-sinistrin (MediBeacon GmbH, #NC1570801) using a miniaturized fluorescence detector (MediBeacon, Germany) attached to the shaved back of the mouse. Before the intravenous injection of 7 mg/100 g FITC-sinistrin, the background fluorescence signal of the skin was recorded for 5 minutes. Fluorescence signals were then continuously recorded for 1-hour post-injection. Data analysis was performed using MPD Studio 2 software (MediBeacon, Germany) after removing the imaging device.

### Behavioral analyses

Open Field Test: To assess general activity and locomotion, mice were tested in an open field arena (45 × 45 cm) with clear Plexiglas walls and a uniformly illuminated white floor. Mice were individually placed at the center and allowed to freely explore for 10 minutes. The mouse’s activity, distance traveled, velocity, and time spent in specific areas of the open field were video-tracked and analyzed using ANYmaze software (Stoelting Co., USA).

Rotarod Test: Whole-body mobility and coordination were assessed using the Rotarod Test. After a 30-minute acclimatization period in the test room, mice were placed on the rod (LE8205, Panlab Harvard Apparatus, Spain), which initially rotated at 4 rpm. The speed was gradually increased from 4 rpm to 40 rpm over 5 minutes, and the latency to fall onto a soft pad was recorded. The test was repeated five times, with a 30-minute interval between trials. After 3 days of training, the latency to fall was calculated based on the average of the 5 trials.

### Selectively depletion of kidney macrophages

Female *Cx3cr1*^CreERT2/+^:*iDTR* mice were first *i.p.* treated with tamoxifen. After that, mice were intravesically given DT on the next consecutive two days. To do so, mice were anaesthetized and intravesically inserted with a PE-10 tubing catheter (RWD Life Science 62324, 0.28 mm 3 OD 0.61 mm) through which 40 ng/g of DT in 80 μl saline was infused. Afterwards, the mice were then placed in a position with their posterior limbs up at a 30-degree angle for 2 hr before wake-up.

### Calcium oxalate kidney stone model

Mice were administered *i.p.* with sodium oxalate (NaOx, Sigma-Aldrich, #71800) at a dose of 60 mg/kg daily for three consecutive days. Some mice were co-treated *i.p.* with a phosphate buffer (Na_2_HPO_4_-NaH_2_PO_4_ with a final concentration of 100 mM phosphate, pH 7.2) at a dose of 100 μl per 25 g of body weight.

### Analyses of CaPi crystals and CaOx stones in the in kidney

Mice were anesthetized and transcardially perfused with cold PBS followed by 4% PFA. The kidneys were then harvested and fixed in 4% PFA overnight, and then dehydrated in 30% sucrose. Large CaOx stones were evaluated by Alizarin Red S (Solarbio, #G1450) staining. For this, frozen sections were incubated in 0.2% Alizarin Red S solution (pH 6.8) at 37 °C for 10 min. For examining CaPi microparticles, mice were *i.v.* infused with 48 nmol/kg OsteoSense 680EX (Revvity, #NEV10020EX) 30 min prior transcardially perfusion. 60-μm-thick slices were prepared for analyses.

### Kidney cortex tubule purification

After perfusion with 4 [ PBS, kidneys were harvested and the cortex of each kidney was separated under a dissecting microscope (Olympus SZ-61-60). Cortex was then mechanically dissociated in RPMI 1640 medium on ice, followed by enzymatic digestion (RPMI 1640 plus with 1 mg/ml Collagenase II (Worthington, #CLS-2), 0.05 mg/ml Soybean Trypsin Inhibitor (Sigma-Aldrich, #T6522-25MG), 1× non-essential amino acid (Corning, #25-025-CIR) and 15 mM HEPES (Sangon Biotech, #E607018-0100)) at 37 [ for 35 min with gentle shaking (∼70 rpm). After centrifugation, cells pellets were resuspended in cold RPMI 1640 medium using an 18G needle and then passed through a 250-μm cell strainer (ThermoFisher Scientific, #87791). Subsequently, tubular fragments, which are larger than 70 μm, were collected using 70-μm cell strainer. The collected proximal tubular fragments were used for RNA extraction and analysis.

### Culture of primary renal tubular epithelial cells

Kidneys from 3-week-old C57BL/6 mice were harvested and immediately placed in DMEM medium (Gibco, #C11995500BT) on ice to maintain cell viability. The tissues were mechanically dissociated and enzymatically digested in 5 ml of DMEM medium containing 1 mg/ml Collagenase II and 5 μl/ml DNase I at 37 °C with gentle shaking for 30 min. Following digestion, the mixture was centrifuged at 300 g for 5 minutes at 4 °C, and the resulting pellet was resuspended in pre-cooled PBS. Tubular fragments which are between 70∼250 μm were collected by passing pellets consecutively through 250-μm and 70-μm cell strainers. The tubule fragments were then resuspended in a complete growth medium consisting of DMEM supplemented with 1% penicillin/streptomycin (Gibco, #15140-122), 10% FBS, 20 ng/ml EGF (PeproTech, #AF-100-15), and 1× ITS (Gibco, #51500056). They were plated in culture dishes containing pre-warmed medium and maintained at 37 °C in a humidified incubator with 5% CO_2_. After 48 hr, the medium was replaced to remove non-adherent cells. Once the epithelial cells reached 70-80% confluence, they were detached, counted, and prepared for further experiments.

### Co-culture primary renal tubular epithelial cells and monocytes

1×10^4^ primary renal tubular epithelial cells were co-cultured with 1×10^5^ monocytes in Transwell system (LABSELECT, #14341) for 24 hr, with epithelial cells in the insert and monocytes in the 24-well plate. Monocytes were used for subsequent RT-PCR.

### Bone marrow-derived macrophages (BMDMs) culture

C57BL/6 mice were euthanized by cervical dislocation. Bone marrow was flushed from both femurs and tibias and cultured in DMEM medium supplemented with 10% FBS, 1% Penicillin-Streptomycin, 2mM L-glutamine (Solarbio, #G0200), 10mM HEPES (Sangon, #E60701) and 15% L929 cell-conditioned medium. On day 6, BMDMs were seeded into wells and prepared for subsequent experiments.

### *In vitro* stimulation with CaPi or CaOx particles

After being washed with PBS, bone marrow-purified monocytes or primary tubular epithelial cells collected from kidney cortex were cultured in DMEM containing 0.1% FBS. Calcium and phosphate concentrations were increased to the indicated final concentrations by adding 1M CaCl_2_ and 1 M phosphate buffer (NaH_2_PO_4_-Na_2_HPO_4_, pH 7.4) to the medium. Monocytes and epithelial cells were cultured in the above medium for 6 hr and 24 hr, respectively, before harvest for RT-PCR analysis.

Some experiments used pre-prepared CaPi particles. To prepare CaPi particles, DMEM was supplemented with CaCl_2_ and phosphate buffer to the final concentrations of 3 mM and 7 mM for calcium and phosphate, respectively. After being gently shaken for 1 hr, the medium was collected and under subject to centrifugation (3,000 g for 15 min). The pellets were resuspended in the same volume of DMEM containing 0.1% FBS, which used for macrophage stimulation. Similarly, to prepare CaOx particles, DMEM was supplemented with CaCl_2_ and NaOx to the final concentrations of 3 mM and 7 mM for calcium and oxalate, respectively.

In some experiments, BMDMs were pre-treated with 5 μM dorsomorphin (MedChemExpress, #HY-13418A), or 20 μM SP600125 (MedChemExpress, #HY-12041) for 30 min and 1 hr, respectively, followed by stimulation with CaPi particles for 2 hr. For chloroquine stimulation, cells were incubated with 100 μM chloroquine (MedChemExpress, #HY-17589A) in cell culture medium for 2 hr.

### LysoTracker and LysoSensor staining

After a 2-hr particle stimulation, cells were washed with PBS, and stained with either 1 μM LysoTracker Green DND-26 (Yeasen, #40738ES50) in PBS which contained 1% FBS for 30 min at 37 °C or 5 μM LysoSensor^TM^ Yellow/Blue DND-160 (PDMPO) (Yeasen, #40768ES50) in PBS for 5 min at 37 °C. LysoTracker was analyzed by flow cytometry on cell basis. For assessing LysoSensor intensity, cells were examined by Synergy HT microplate reader (BioTek, Synergy H1) with fluorescence emission at 440 nm (excitation 329 nm) and emission at 540 nm (excitation 384 nm). An emission-intensity ratios (I_440_/I_540_) was used to quantify changes in lysosomal pH.

### Western Blot analysis

Cells were collected and lysed in RIPA Lysis Buffer (Beyotime Biotechnology, #P0013B) with a cocktail of protease inhibitors (Thermo Scientific, #A32965). After centrifugation at 12,000 g for 15 min at 4 [, supernatants were collected for Western blotting according to a standard procedure. Antibodies against Phospho-AMPKα (Thr172) (Cell Signaling Technology, #2535, 1:1,000), AMPKα (Cell Signaling Technology, #2532, 1:500) were used, and an antibody against β-actin was used as a loading control (ABclonal, #AC026, 1:10,000). Chemiluminescence was measured on a ChemiScope3300 imaging system (Clinx).

### Statistics

Statistical analysis was performed with GraphPad Prism 6.0 (GraphPad). Data were presented as mean±SEM. Two-tailed one-way ANOVA and two-way ANOVA with Tukey’s multiple-comparisons testing was used. Student’s *t* tests were used to compare two groups. *P*<0.05 were considered significant.

