## Supplementary Figures for "Kidney cortex macrophages strategically reabsorb phosphate from the urine to prevent the formation of mineral stones"

**Supplementary Figures and Legends**

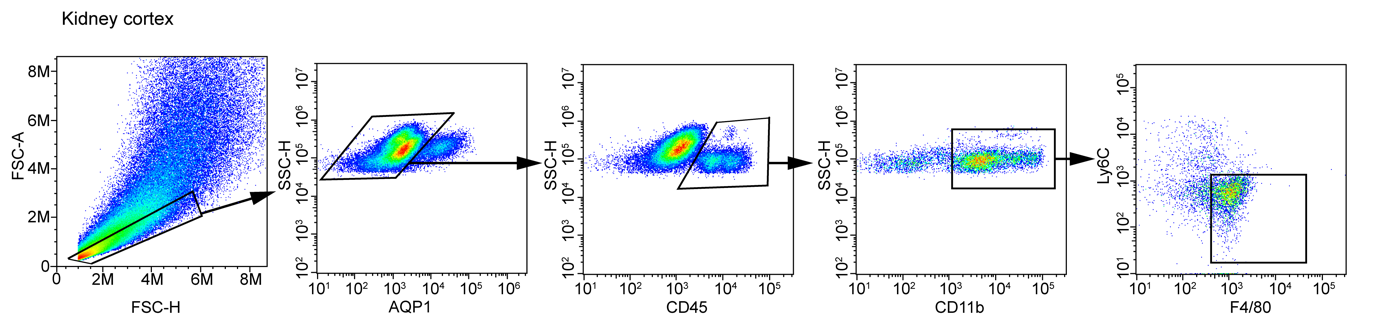
**Figure S1**

**Figure S1 Gating strategy for kidney cortex macrophages.**

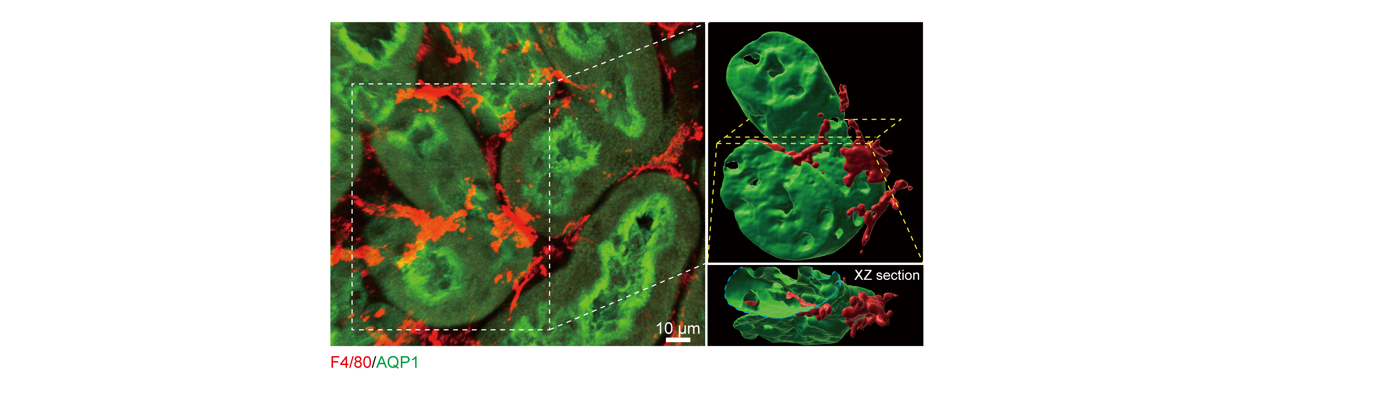
**Figure S2**

**Figure S2** Confocal image of the spatial relationship between AQP1^+^ epithelium and F4/80^+^ cell protrusions. The magnifications show the corresponding 3D reconstructions. Z-projections of 22.5 μm.

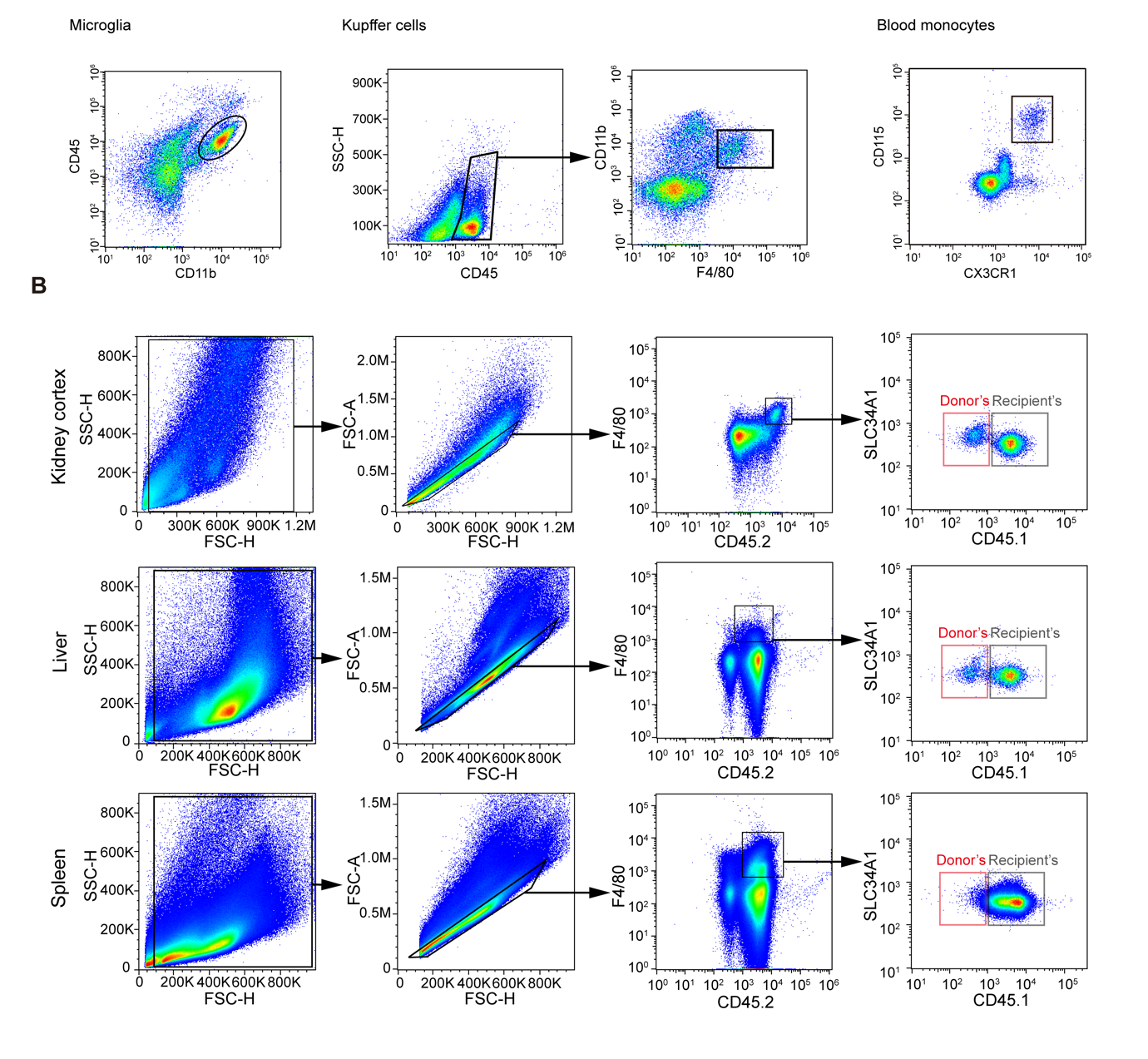
**Figure S3**

**Figure S3 Gating strategies.** (**A**) For microglia from the brain, Kupffer cells from the liver and monocytes from the blood. (**B**) For distinguishing donor-derived MØ (CD45.2) from recipients’ own MØ (CD45.1/CD45.2) in kidney cortex, liver and spleen.

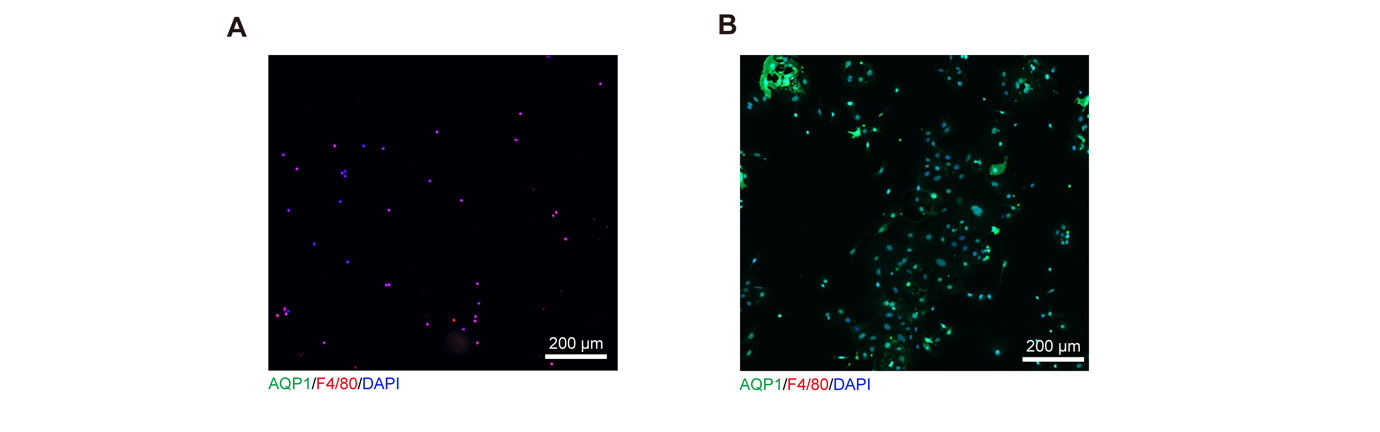
**Figure S4**

**Figure S4 The purity of the purified cells from kidney cortex.** (**A**) Following the protocol of kidney cortex MØ purification, cells were seeded into the culture dish and they were then under F4/80 and AQP1 staining. (**B**) Following the protocol of kidney cortex tubule purification, cells were seeded into the culture dish and they were then under F4/80 and AQP1 staining.

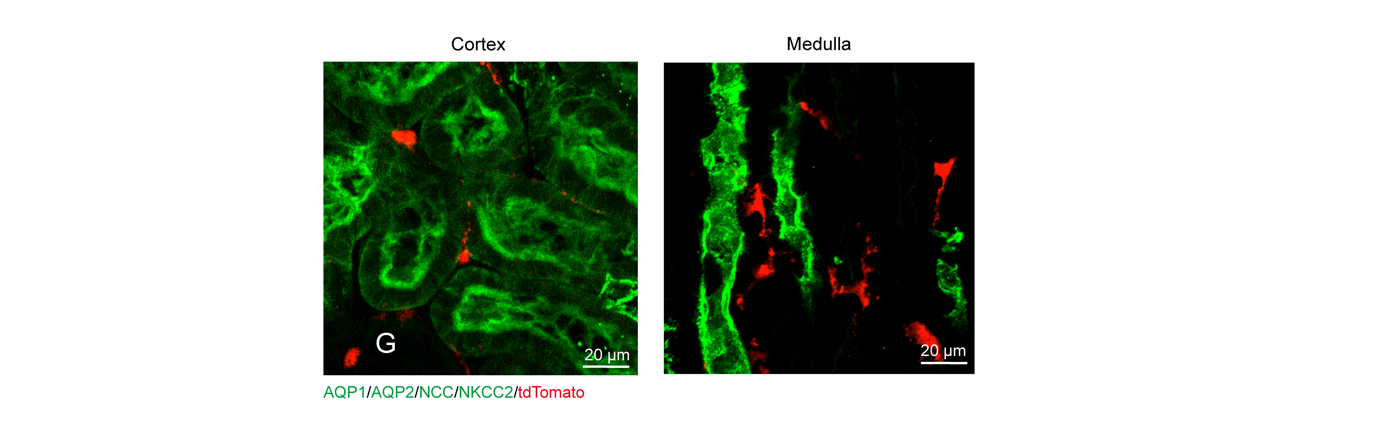
**Figure S5**

**Figure S5 Kidney tubule epithelial cells do not express CX3CR1.** CX3CR1 expression in the cortex and medulla of kidneys derived from *Cx3cr1*^CreERT2/+^:Ai14 mice were examined. AQP1, AQP2, NCC and NKCC2 together mark most renal tubules. G, glomerulus. Tubule epithelial cells (green) do not express tdTomato. The tdTomato^+^ cells were macrophages, as shown before (Ref: 20)

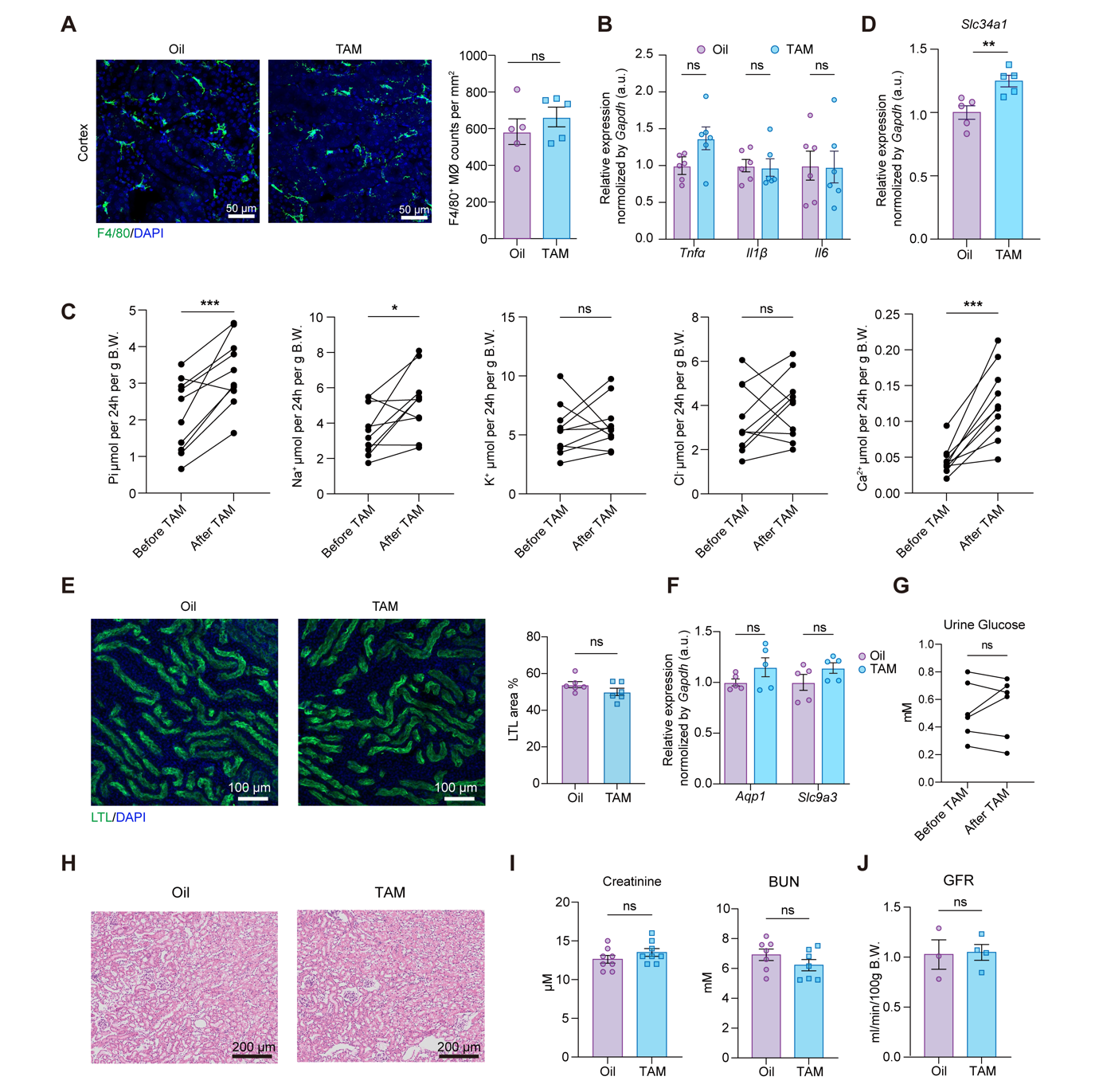
**Figure S6**

**Figure S6 Phenotypes of mice abrogated with *Slc34a1* expression in kidney cortex macrophages.** These experiments followed the experimental scheme shown in Fig.2A. (**A**) Representative images of cortex sections and statictics show the densities of F4/80^+^ MØ in *Slc34a1*^ΔMØ^ mice treated with corn oil (Oil) or Tamoxifen (TAM). (**B**) The purified cortex MØ were tested for the expression of the indicated pro-inflammatory cytokine. (**C**) Urine output of the indicated electrolytes adjusted by body weight before and after TAM treatment. (**D**) The expression of *Slc34a1* in purified tubules from kidney cortex at the end of the experimental scheme. (**E**) The density and morphology of LTL^+^ cortical proximal tubules at the end of the experimental scheme. Each dot indicates the average value of eight 639 × 639 μm^2^ FOVs from one mouse. (**F**) The expression of *Aqp1* and *Slc9a3* in purified tubules from kidney cortex at the end of the experimental scheme. (**G**) Urine glucose concentrations before and after TAM treatment. (**H**) Representative H&E staining shows the general architecture of the kidneys under the indicated treatment. (**I**) Blood urea nitrogen (BUN) and creatine concentrations, and (**J**) Glomerulus filtration rate (GFR) at the end of the experimental scheme. ns, not significant. **P*<0.05, ***P*<0.01, ****P*<0.005, two-tailed unpaired *t* test (A, B, D, E, F, I, J), and by two-tailed paired *t* test (C, G). Data are depicted as mean±SEM. Data are derived from at least 2 independent experiments.

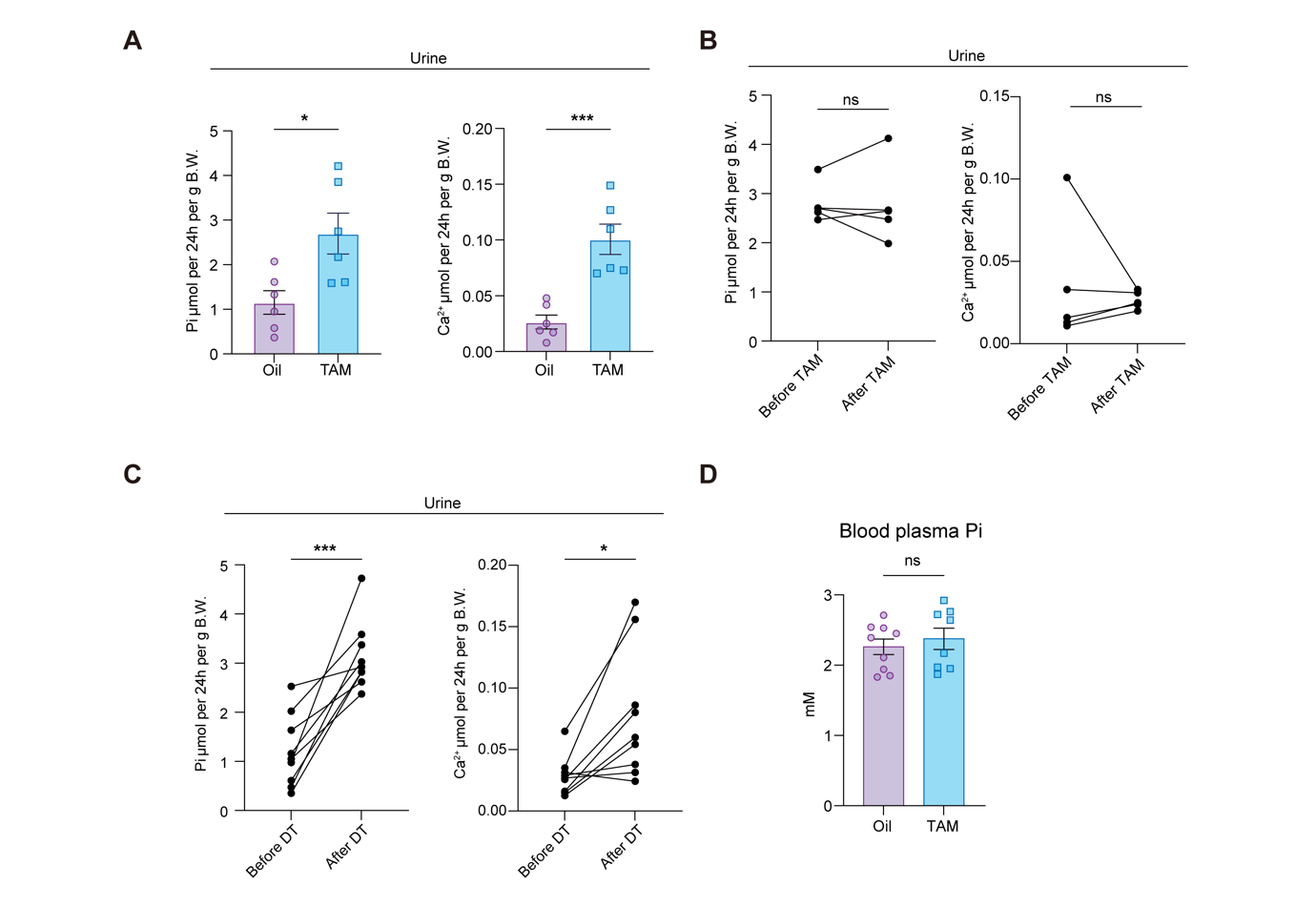
**Figure S7**

**Figure S7 Pi and calcium concentrations in the urine and Pi concentration in the blood.** (**A**) Both kidney of *Slc34a1*^ΔMØ^ mice were subjected to subcapsular administration of corn oil or tamoxifen. Urine Pi and calcium output was measured. (**B**) Before and after tamoxifen *i.p.* treatment, urine Pi and calcium output of C57BL/6 was measured. (**C**) Before and 4 d after intravesical application of DT to *Cx3cr1*^CreERT2/+^:*iDTR* mice, urine Pi and calcium output was measured. (**D**) Pi concentration in the blood plasma of *Slc34a1*^ΔMØ^ mice on day 8 after corn oil or tamoxifen treatment. ns, not significant. **P*<0.05, ****P*<0.005 by two-tailed unpaired *t* test (A, D) and by two-tailed paired *t* test (B, C). Data are depicted as mean±SEM. Data are derived from 2 independent experiments.

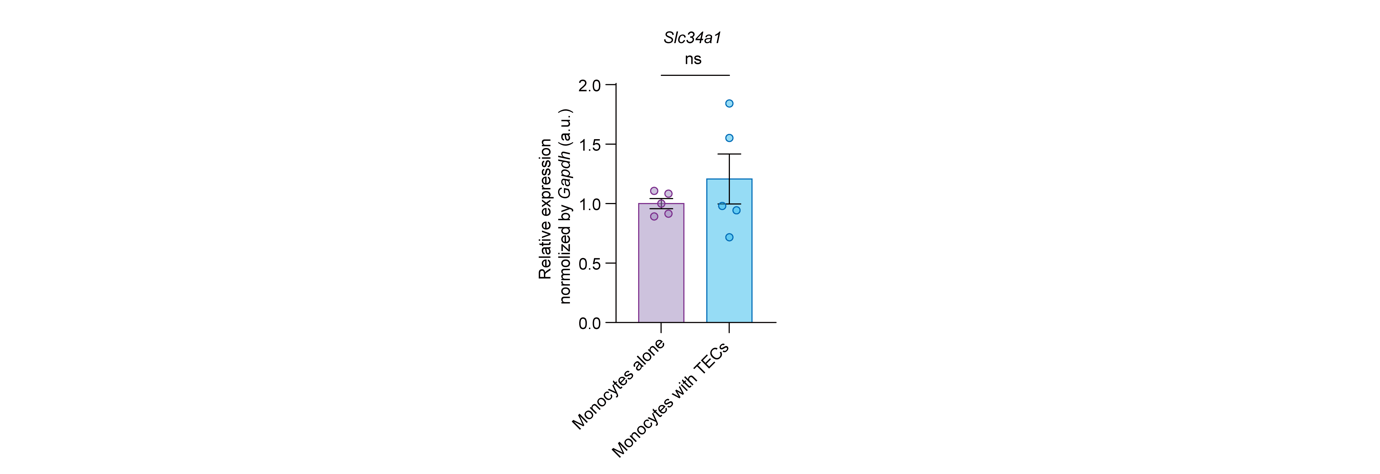
**Figure S8**

**Figure S8 Co-culture of monocytes with tubular epithelial cells collected from kidney cortex.** The purified cortex tubules were first cultured for 48 hr to be monolayer epithelial cells (TECs). Then, bone marrow-purified monocytes were cultured alone or with TECs for 24 hr in a transwell system. *Slc34a1* expression in monocytes were evaluated by RT-PCR. ns, not significant by two-tailed unpaired *t* test. Data are depicted as mean±SEM. Data are derived from 5 independent experiments.
